# Kinetic Properties of Microbial Exoenzymes Vary with Soil Depth but Have Similar Temperature Sensitivities Through the Soil Profile

**DOI:** 10.1101/2021.07.02.450965

**Authors:** Ricardo J. Eloy Alves, Ileana A. Callejas, Gianna L. Marschmann, Maria Mooshammer, Hans W. Singh, Bizuayehu Whitney, Margaret S. Torn, Eoin L. Brodie

## Abstract

Current knowledge of the mechanisms and responses of soil organic matter (SOM) turnover to warming is mainly limited to surface soils, although over 50% of global soil carbon is contained in subsoils. Deep soils have different physicochemical properties, nutrient inputs and microbiomes, which may harbor distinct functional traits and lead to different SOM dynamics and temperature responses. We hypothesized that kinetic and thermal properties of microbial exoenzymes, which mediate SOM depolymerization, vary with soil depth, reflecting microbial adaptation to distinct substrate and temperature regimes. We determined the Michaelis-Menten (MM) kinetics of three ubiquitous enzymes involved in carbon (C), nitrogen (N) and phosphorus (P) acquisition at six soil depths down to 90 cm at a temperate coniferous forest, and their temperature sensitivity based on Arrhenius and Macromolecular Rate Theory (MMRT) models over six temperatures between 4-50°C. Maximal enzyme velocity (*V*_max_) decreased strongly with depth for all enzymes, both on a dry soil mass and a microbial biomass C basis, whereas their affinities increased, indicating adaptation to lower substrate availability. Surprisingly, microbial biomass-specific catalytic efficiencies also decreased with depth, except for the P-acquiring enzyme, indicating distinct nutrient demands at depth relative to microbial abundance. These results indicated that deep soil microbiomes encode enzymes with intrinsically lower turnover and/or produce less enzymes per cell, likely reflecting distinct life strategies. The relative kinetics between different enzymes also varied with depth, suggesting an increase in relative P demand with depth, or that phosphatases may be involved in C acquisition. Warming consistently led to increased *V*_max_ and catalytic efficiency of all enzymes, and thus to overall higher SOM-decomposition potential, but enzyme temperature sensitivity was similar through the soil profile based on both Arrhenius/Q_10_ and MMRT models. Nevertheless, temperature directly affected the kinetic properties of different enzyme types in a depth-dependent manner, and thus the relative depolymerization potential of different compounds. Our results indicate that kinetic and thermal properties of exoenzymes are intrinsic traits of soil microbiomes adapted to distinct physicochemical conditions associated with different soil depths, and improve our conceptual understanding of critical mechanisms underlying SOM dynamics and responses to warming through the soil profile.

## Introduction

Soils are estimated to contain ∼3,000 Gt carbon (C), which is more than all C in the atmosphere and in living biomass combined (Köchy et al., 2015). The dynamics of the large soil C reservoir is sensitive to climate change, and C losses as carbon dioxide (CO_2_) are expected to become a major positive feedback to global warming through increased soil organic matter (SOM) decomposition (Crowther et al., 2016; Van Gestel et al., 2018). An estimated 55 ± 50 Gt C may be lost globally from just 1°C warming of the upper 10 cm of soil alone (Crowther et al., 2016). While current model predictions of C dynamics and responses to climate change are largely based on surface soils (Crowther et al., 2016; Trumbore, 2009; Van Gestel et al., 2018), soils below 20 cm contain up to 50% of the global soil C budget within the top 1 m of soil (Balesdent et al., 2018; Jobbágy and Jackson, 2000). These subsoils are predicted to warm at rates similar to those of surface soils (Soong et al., 2020), and recent *in situ* deep soil warming experiments have shown uniform warming responses down to 100-120 cm depth leading to soil C losses at least three times higher than those estimated based on surface soils alone (Hanson et al., 2020; Hicks Pries et al., 2017; Nottingham et al., 2020; Soong et al., 2021). Despite these observations, relatively little is known about the microbial mechanisms and interactions mediating SOM turnover and CO_2_ emissions, and their responses to environmental changes in subsoils (Gross and Harrison, 2019; Rumpel and Kögel-Knabner, 2011), which are essential to improve predictions of SOM dynamics in response to warming.

Soil physicochemical properties and environmental conditions, such as nutrient inputs, temperature, moisture, mineralogy, and organic matter composition vary markedly with depth, creating distinct environments for the microbial processes that mediate SOM transformations (Blume et al., 2002; Fierer et al., 2003a; Jones et al., 2018; Rumpel and Kögel-Knabner, 2011; Salomé et al., 2010; Trumbore, 2000). The rate-limiting steps in SOM decomposition are primarily catalyzed by microbial exoenzymes, which depolymerize plant and microbial residues into lower molecular weight compounds that are assimilated by both plants and microbes (Burns et al., 2013; Davidson and Janssens, 2006). The kinetic and thermal properties of exoenzymes are therefore fundamental determinants of SOM turnover, nutrient availability, soil C stability, and greenhouse gas emissions, as well as their responses to environmental changes (Chen et al., 2018; Davidson and Janssens, 2006; Sinsabaugh and Shah, 2012; Wallenstein et al., 2011). In addition to the large diversity of exoenzymes targeting different organic compounds, evolutionarily distinct exoenzymes that catalyze the same reactions (i.e., isozymes) can vary widely in their kinetic properties, namely their maximal substrate turnover (*K*_cat_) and related maximal reaction velocity (*V*_max_), their Michaelis constant (*K*_m_), which is inversely proportional to their affinity, and their catalytic efficiency (*K*_cat_*/K*_m_) (Khalili et al., 2011; Nannipieri et al., 2012; Sinsabaugh and Shah, 2012; Tischer et al., 2015). These properties constitute microbial evolutionary adaptations and trade-offs related to resource supply and demand, associated with distinct ecological niches (Allison et al., 2011; Ho et al., 2017; Malik et al., 2020; Sinsabaugh and Shah, 2012). Temperature is also a major factor controlling microbial community assembly, growth and functionality (Allison and Treseder, 2008; Bradford, 2013; Cavicchioli et al., 2019; Lax et al., 2020), and warming has been shown to change the abundance of diverse taxa and functional groups through the soil profile (Dove et al., 2021; Jiang et al., 2020). Moreover, optimal enzyme temperatures are broadly correlated with the optimal growth temperatures of their organisms, as well as with the frequency of specific metabolic pathways, reflecting a concerted evolutionary adaptation to temperature and associated selective pressures (Engqvist, 2018; Somero, 2004). Therefore, variation in substrate and temperature regimes through the soil profile is likely to select for microbiomes producing enzymes with distinct kinetic and thermal properties, which may impose depth-dependent constraints on SOM turnover and responses to warming (Allison and Treseder, 2008; Carrillo et al., 2018; Cavicchioli et al., 2019; Isobe et al., 2019; Nunan et al., 2020; Xu et al., 2021). Microbial community composition and functional potential have indeed been shown to vary strongly with soil depth, reflecting selective adaptation to distinct niches (Blume et al., 2002; Brewer et al., 2019; Diamond et al., 2019; Dove et al., 2021; Eilers et al., 2012; Fierer et al., 2003b; Hansel et al., 2008; Hartmann et al., 2009; Jiao et al., 2018; Liu et al., 2019; Turner et al., 2017; Yan et al., 2019; Zosso et al., 2021). At the same time, exoenzyme activities in nature are dependent on multiple factors that can directly or indirectly modulate their kinetics, thermodynamics, and expression, beyond the intrinsic traits of the microbiome and the enzymes they encode. In particular, microbe-plant interactions, soil properties, and environmental conditions all affect enzyme expression, turnover, mobility, and substrate accessibility (Bradford, 2013; Burns et al., 2013; Davidson and Janssens, 2006; Tang and Riley, 2019; Wallenstein et al., 2011). Consequently, the effective kinetics of mixed exoenzyme pools in complex environments are emergent properties that reflect not only the summation of traits from distinct isozymes and organisms, but also direct and indirect interactions between enzymes and the environment (Burns et al., 2013; Davidson and Janssens, 2006; Sinsabaugh and Shah, 2012).

Soil exoenzyme activities and their environmental controls have been extensively studied in the context of warming and other environmental changes, as indicators of SOM decomposition activity and nutrient availability, but have largely focused on one kinetic property (i.e., *V*_max_) and on surface soils (Allison et al., 2011; Burns et al., 2013; Sinsabaugh and Shah, 2012). Given their critical role in soil C stability and CO_2_ emissions, several efforts have been made to integrate kinetic and thermal properties of exoenzymes to improve the mechanistic understanding of SOM dynamics in response to warming (Allison et al., 2018; Alster et al., 2016a, 2020; Burns et al., 2013; German et al., 2012; Loeppmann et al., 2016a; Razavi et al., 2015, 2016; Wallenstein et al., 2011). Several studies have investigated exoenzyme activities through the soil profile (Darby et al., 2020; Dove et al., 2020; Gelsomino and Azzellino, 2011; Jing et al., 2017; Kramer et al., 2013; Loeppmann et al., 2016a; Schnecker et al., 2014, 2015; Stone et al., 2014; Taylor et al., 2002; Venkatesan and Senthurpandian, 2006). However, nearly all depth-resolved soil studies, possibly with just one exception (Loeppmann et al., 2016a), have relied on potential enzyme activity assays based on single substrate concentrations, and have not experimentally determined the Michaelis-Menten (MM) kinetics required to estimate the true enzyme *V*_max_, as well as *K*_m_ and catalytic efficiency (in the context of mixed enzyme pools and complex environments; Wallenstein et al., 2011). Moreover, the temperature sensitivity of soil exoenzymes over the whole soil profile has rarely been characterized (Steinweg et al., 2013), and studies that determined both the MM kinetics of exoenzymes and their direct temperature sensitivity are scarce, even for surface soils (Allison et al., 2018; Razavi et al., 2015, 2016).

The temperature sensitivity of soil exoenzymes and other biogeochemical processes has been typically determined based on the linear Arrhenius model and related Q_10_ coefficient, which represents a simple empirical metric expressing variation in activity rates at every 10°C change in temperature (Alster et al., 2020). However, it has been argued that the Q_10_ coefficient may not reliably represent soil biological processes, as it lacks a biological and mechanistic basis, and does not capture the unimodality of typical enzyme reactions (Alster et al., 2020; Hobbs et al., 2013). These caveats possibly explain the frequent inability of Q_10_ values to describe observed temperature responses of soil biological processes, and lack of comparability between studies (Alster et al., 2020). Macromolecular Rate Theory (MMRT) has been recently proposed as a more realistic model of enzyme temperature sensitivity based on thermodynamics and the change in heat capacity associated with enzyme catalysis, which accounts for declines in enzyme activity below thermal denaturation temperatures (Hobbs et al., 2013). MMRT can thus appropriately capture the unimodal behavior of enzyme response to temperature, and describes temperature sensitivity as comprising three fundamental components: temperature optimum (*T*_opt_), the temperature at which reaction rates are maximal; point of maximum temperature sensitivity (TS_max_), the temperature at which reaction rates change the most; and change in heat capacity (Δ*C*_p_^‡^), which describes the degree of curvature of the parabolic response of reaction rates to temperature (Alster et al., 2020). A limited number of studies have applied MMRT to soil biological activities, including exoenzymes in soils and cultures of soil microbes, where it could describe temperature responses more coherently than Arrhenius models and provide more realistic interpretations of temperature sensitivity (Alster et al., 2016a, 2016b, 2018; Robinson et al., 2017; Schipper et al., 2014). However, to our knowledge, MMRT has never been used to investigate the temperature sensitivity of exoenzymes over the whole soil profile.

Different soil models have been developed to represent exoenzyme kinetics, thermodynamics, ecological stoichiometry, enzyme diffusion, and interactions with environmental factors (German et al., 2012; Sinsabaugh and Shah, 2012; Sulman et al., 2014; Tang and Riley, 2015, 2019; Wang et al., 2015; Wang and Allison, 2019; Wieder et al., 2014). However, these processes have only recently started to be incorporated into depth-resolved soil biogeochemical models (Dwivedi et al., 2019; Wang et al., 2021), are rarely considered in fully coupled ecosystem scale models (Grant, 2013; Pasut et al., 2021), and are entirely unrepresented in current Earth system models. Moreover, exoenzyme kinetics, when included in depth-resolved models, are represented as a function of microbial biomass, and not as explicit properties that may vary independently due to differences in microbial life strategies or microbe-substrate interactions.

We investigated how kinetic properties and temperature sensitivity of microbial exoenzymes vary with soil depth, and if they may represent depth-dependent traits associated with microbiomes adapted to distinct soil environments. We hypothesized that: (i) enzyme *V*_max_ declines with depth, in concert with declines in substrate concentrations and overall nutrient demand; (ii) enzyme affinities and catalytic efficiencies increase with depth to maximize resource acquisition under low substrate concentrations; (iii) variation of kinetic properties with depth differs between C-, N- and P-acquiring enzymes, reflecting differences in relative substrate availability and demand; (iv) temperature sensitivity of exoenzymes increases with depth, reflecting selection of enzymes adapted to lower and narrower temperature ranges in deeper soils. We determined the MM kinetics and catalytic efficiencies of the hydrolytic enzymes β-glucosidase (BG), leucine aminopeptidase (LAP) and acid phosphatase (AP) (involved in C, N and P acquisition, respectively), as a function of both soil dry mass and microbial biomass C, in soils collected at six depths down to 90 cm at a temperate coniferous forest. Furthermore, we investigated enzyme temperature sensitivity based on the Arrhenius model and Q_10_ coefficients, and on the MMRT model over six temperatures between 4-50°C, following a fully factorial experimental design considering substrate type and concentration, soil depth and temperature.

## Materials and Methods

### Site Description and Sample Collection

Soil samples were collected at the University of California Blodgett Experimental Forest, Sierra Nevada, CA (120°39’40” W; 38°54’43” N), described by Hicks Pries et al. (2018). Briefly, Blodgett forest is located in a Mediterranean climate with mean annual precipitation of 1,660 mm and a mean annual air temperature of 12.5°C. The soil was classified as Alfisol of granitic origin, and has a developed O horizon. The site is a mixed coniferous forest with ponderosa pine (*Pinus ponderosa*), sugar pine (*Pinus lambertiana*), incense cedar (*Calodefrus decurrens*), white fir (*Abies concolor*) and douglas fir (*Pseudotsuga menziesii*) as dominant tree species. The mean annual soil temperature ranges between 11.5 and 10.4°C at 5 and 100 cm depths, respectively, although soil temperatures vary annually between 0-29°C, 1-19°C and 2-16°C at 5, 30 and 100 cm depth, respectively. Three soil cores were collected in July 2019 using a 4.78 cm diameter soil corer with a 10 kg hand-held slide-hammer. The surface litter layer of the O horizon was removed prior to sampling, and mineral soil samples were recovered sequentially in 10 cm increments down to 90 cm depth. Samples were kept cold during transportation to the laboratory, were they were sieved to 2 mm and stored at 4°C. Samples were analyzed within approximately a week of collection. To ensure the accessibility and discoverability of the samples generated here, and to align with the National Science Foundation’s guidelines of effective data practices, all samples have been registered with IGSN Global Sample Numbers through the System for Earth Sample Registration (SESAR). SESAR is maintained by the GeoInformatics Research Group of the Lamont-Doherty Earth Observatory at https://www.geosamples.org/. Sample IGSNs are shown in Table S1.

### Exoenzyme Activity Assays

Extracellular hydrolytic enzyme activities were determined fluorometrically according to standard assays (German et al., 2011b) at six depth intervals, following the experimental design in Table 1. Briefly, we used the methylumbelliferone (MUF)-linked substrates MUF-β-D-glucopyranoside and MUF-phosphate for determination of β-glucosidase (BG) and acid phosphatase (AP) activities, respectively. Leucine aminopeptidase (LAP) activity was determined using the substrate L-leucine-7-amido-4methylcoumarin (AMC). Assays were performed for each of six soil depths from each of three replicate soil cores, by combining 200 µL of soil homogenate with 50 µL of fluorogenic substrate solution in each microplate well. Soil homogenates were prepared using 1 g of fresh soil in 100 mL 50 mM acetate buffer with pH 5.5. Substrate solutions, soil homogenates, serial dilutions of standards in the absence or presence of soil homogenate (quenching controls), and blank controls in the absence or presence of each of the eight substrate concentrations were prepared in the same buffer. Eight serial dilutions of MUF and AMC standards, blank controls with only added substrates, and quenching controls with soil homogenates, but no MUF or AMC added, were performed in duplicate. Each enzyme was assayed individually over a range of eight substrate concentrations, as follows: 10, 30, 60, 100, 150, 250, 450 and 800 µM for BG; 10, 20, 40, 70, 110, 190, 350 and 600 µM for LAP; and 10, 40, 80, 130, 200, 350, 700 and 1200 µM for AP (Table 1). Parallel assays for each sample, enzyme and substrate concentration were incubated at 4, 10, 16, 25, 35 or 50°C, and fluorescence was recorded after approximately 1, 4 and 24 h to determine the optimal incubation time. Four analytical replicates were measured per sample for each combination of enzyme, substrate concentration and temperature. Incubation temperatures were selected in order to capture the unimodal response predicted by MMRT with *T*_opt_ values well above native temperatures, as observed by previous studies of exoenzymes from temperate environments (Alster et al., 2016b), while including the temperature range and approximate seasonal averages at our experimental site.

**Table 1.** Experimental set-up of enzyme potential activity assays.

| Enzyme | Fluorogenic substrate | Substrate concentration (μM) | Soil depth (cm) | Temperature (°C) | Incubation time (h) |
| --- | --- | --- | --- | --- | --- |
| β-glucosidase (BG)<br>EC 3.2.1.21 | 4-methylumbelliferyl-β-D-glucopyranoside | 10, 30, 60, 100, 150, 250, 450, 800 | 0-10 | 4 | 2 |
|  |  |  | 10-20 | 10 | 4 |
| Leucine aminopeptidase (LAP)<br>EC 3.4.11.1 | L-leucine-7-amido-4-methylcoumarin | 10, 20, 40, 70, 110, 190, 350, 600 | 30-40 | 16 | 24 |
|  |  |  | 50-60 | 25 |  |
| Acid Phosphatase (AP)<br>EC 3.1.3.2 | 4-methylumbelliferyl phosphate | 10, 40, 80, 130, 200, 350, 700, 1200 | 60-70 | 35 |  |
|  |  |  | 80-90 | 50 |  |

### Microbial Biomass C and Dissolved C and N Pools

Microbial biomass C (MBC), dissolved organic C (DOC) and total dissolved N (TDN) were determined at every 10 cm depth interval between 0-90 cm depth. MBC was estimated using the chloroform-fumigation extraction method (Brookes et al., 1985). Five-gram soil samples were fumigated in 50 mL closed vials containing a jumbo cotton ball soaked with ethanol-free chloroform over, but not touching, the soil, for 7 days, with chloroform replenished on day 4. Fumigated and a non-fumigated soil samples were extracted with 25 mL of 0.5 M K_2_SO_4_ on an orbital shaker table for 60 min, then gravity filtered through pre-leached #42 Whatman filter paper, and frozen until further analysis. DOC and TDN in fumigated and non-fumigated samples were quantified using a Lotix Combustion TOC/TN Analyzer (Teledyne Tekmar, Mason, Ohio, USA). No correction factor (k_EC_) was applied to account for incomplete microbial biomass lysis during the fumigation.

### Data Analyses

All data manipulations and analyses were performed in R versions 3.6.1-4.0.4 (R Core Team, 2020). Net fluorescence in the enzyme assays, including quenching corrections, were calculated following German et al., (2011b). Analytical outliers were identified based on the Interquartile Range method, and a maximum of one value was excluded out of the four analytical replicates per sample. Negative values due to analytical error were excluded from the dataset. Enzyme maximal velocity (*V*_max_) and Michaelis constants (*K*_m_) were computed by fitting a 2-parameter Michaelis-Menten (MM) model over all analytical replicates per substrate concentration using the *drm* function in the *drc* package (Ritz et al., 2015), with a data-driven self-starter function specific to the model. Following preliminary analyses, and when necessary, we excluded data points corresponding to one of the eight individual substrate concentrations for which all analytical replicates consistently did not fit the distribution of the remaining data (i.e., due to technical errors during assay preparation). In order to alleviate variance heterogeneity of analytical replicates between substrate concentrations, we applied a Box-Cox transformation to all models using the *boxcox* function in the *drc* package (Ritz et al., 2015). Comparison between the parameters *V*_max_ and *K*_m_ estimated based on transformed and non-transformed models showed that Box-Cox transformation improved the fit of models with substantial analytical variance, but had a marginal or no effect on parameters estimated by models with initial good fit. Individual models yielding nonsignificant *V*_max_ or *K*_m_ estimates (*p* > 0.05) after Box-Cox transformation were considered to have bad fit and were thus excluded from further analyses (excluded 11 out of 324 models). To determine the optimal assay incubation time at each temperature, we compared MM models fit to data collected after each of three sequential incubation periods (1, 4 and 24 h). We selected the minimum incubation period necessary to reach the highest *V*_max_ value, under the assumptions that lower *V*_max_ values reflected either insufficient incubation time for reactions to reach saturation, or loss of fluorescence due to prolonged incubation after saturation had been reached. The same incubation period was consistently selected for all assays performed at the same temperature. *V*_max_ was expressed per mass of dry soil as *V*_max/ds_ (nmol g^-1^ h^-1^) and per unit of microbial biomass C (MBC) as *V*_max/MBC_ (nmol µg MBC^-1^ h^-1^). The apparent catalytic efficiency (CE_ds_) was calculated as:

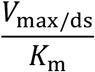

and the biomass-specific catalytic efficiency (CE_MBC_) as:

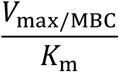

Q_10_ coefficients were calculated over the full experimental temperature range (six temperatures from 4 to 50°C) and over a realistic field range (five temperatures from 4 to 35°C) following the approach by Allison et al. (2018). Briefly, the degree of change in *V*_max_, *K*_m_ or CE per °C was inferred based on linear regressions between the natural logarithm of each parameter and temperature, and converted to Q_10_ values based on the relationship:

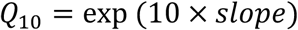

The Arrhenius activation energy (*E*_a_) was calculated based on the slope of the linear regression between ln(*V*_max_) and 1/*T*, and the relationship:

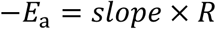

where *T* is the temperature in kelvin and *R* is the universal gas constant. Linear regression models were calculated using the *lm* function in the *stats* package native to R (R Core Team, 2020). The change in heat capacity (Δ*C*_p_^‡^), temperature optimum (*T*_opt_) and point of maximum temperature sensitivity (TS_max_) were calculated by fitting ln(*V*_max_) over the six incubation temperatures between 4-50°C using the Macromolecular Rate Theory (MMRT) model, according to the equations and definitions described by Alster et al. (2020). The reference temperature *T*_0_ was set to 315 K to best match the measured data, following the recommendations by Alster et al. (2020). Model fit comparisons were based on the Akaike Information Criterion (AIC) and respective relative likelihoods, corrected AIC (AICc), and Bayesian Information Criterion (BIC), following the guidelines by Burnham and Anderson, (2004). AIC, BIC, and adjusted R^2^ values of the linear models were extracted from the linear regression model computed with the *lm* function in the *stats* package (R Core Team, 2020). AICc of all models, and AIC and BIC of the nonlinear models were calculated using the R package *AICcmodavg* (Mazerolle, 2020). One-way and two-way Analyses of Variance (ANOVA) were performed with the *aov* function, followed by post-hoc Tukey’s tests using the function *TukeyHSD* with *p* values adjusted for multiple comparisons, using the *stats* package (R Core Team, 2020). Compact letter displays for the Tukey’s tests were computed with the function *HSD*.*test* in the package *agricolae* (de Mendiburu and Yaseen, 2020). Assumptions of ANOVA were tested based on Levene’s tests with the *leveneTest* function in the package *car* (Fox and Weisberg, 2019), Shapiro-Wilk tests with the *shapiro*.*test* function in the package *stats* (R Core Team, 2020), skewness of residuals with the *skewness* function in the package *agricolae* (de Mendiburu and Yaseen, 2020), and plots of homogeneity of residuals’ variance and normality of residuals (Q-Q plots). Data was ln-transformed as necessary, and all tests reported as significant were based on a *p* value < 0.05. Figure displays were prepared with the package *cowplot* (Wilke, 2020). The maximum percentage of variation (i.e., decline) in kinetic parameters with depth, per temperature, was calculated as the percentage of difference between the highest and lowest values within the upper and lower depth intervals mentioned in the text, for example:

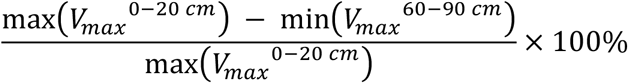

As the ANOVA showed that variation in kinetic parameters with depth was not dependent on temperature, the percentages of variation with depth are reported as the average decline among all temperatures, per enzyme and kinetic parameter. The inconsistently high mean *K*_m_ values only at 16°C was excluded from those calculations.

## Results

### Exoenzyme kinetics vary with soil depth

We determined the MM kinetics of the enzymes acid phosphatase (AP), β-glucosidase (BG), and leucine aminopeptidase (LAP) in soils collected at six depth intervals from triplicate soil cores down to 90 cm (0-10, 10-20, 30-40, 50-60, 60-70 and 80-90 cm), at six temperatures between 4 and 50°C (4, 10, 16, 25, 35 or 50°C) (Table 1). Activity of the three enzymes showed typical MM behavior, and the kinetic parameters *V*_max_ and *K*_m_ were computed by fitting a nonlinear regression model of the MM equation to potential enzyme activities over eight substrate concentrations with four analytical replicates. Variation in kinetic parameters with depth and interactions with temperature were tested using two-way fixed effects Analysis of Variance (ANOVA) with depth and temperature as independent factors, followed by post-hoc Tukey’s tests.

The *V*_max_ of all three enzymes, estimated on a dry soil mass basis (*V*_max/ds_), declined significantly with soil depth over all temperatures (*p* < 0.001), and differences among depths were not dependent on temperature (i.e., no significant depth × temperature interaction) (Figure 1A, Table 2). Mean *V*_max/ds_ declined almost continuously from the soil surface (0-20 cm) down to 60-90 cm by up to 96.4 ± 0.4% (mean ± standard error; see Materials and Methods for details) across all enzymes and temperatures. This variation was only significant between three to four depth ranges, which differed between enzymes (Figure 1A, Table S2): *V*_max/ds_ of BG declined progressively down to 60 cm, but not below that depth; *V*_max/ds_ of AP declined only over the mid-depth range, from 20 to 30 cm and from 40 to 60 cm; *V*_max/ds_ of LAP also did not vary within the upper 20 cm, but declined gradually down to a lower depth than that of AP, namely from 20 to 30 cm, from 40 to 50 cm and from 60 to 80 cm.

**Table 2.** Two-way fixed effects ANOVA of kinetic parameters with depth, with depth and temperature as independent factors. Differences were considered significant at *p* < 0.05

| Enzyme | Factor | $V_{\max}/ds$ | | | $V_{\max}/MBC$ | | | $K_m$ | | | $CE_{ds}$ | | | $CE_{MBC}$ | | |
| --- | --- | --- | --- | --- | --- | --- | --- | --- | --- | --- | --- | --- | --- | --- | --- | --- |
|  |  | Df | F value | p value | Df | F value | p value | Df | F value | p value | Df | F value | p value | Df | F value | p value |
| BG | Depth | 5 | 89.32 | < 0.05 | 5 | 100.95 | < 0.05 | 5 | 30.03 | < 0.05 | 5 | 45.42 | < 0.05 | 5 | 27.37 | < 0.05 |
|  | Temperature | 5 | 19.94 | < 0.05 | 5 | 54.16 | < 0.05 | 5 | 2.64 | 0.03 | 5 | 14.55 | < 0.05 | 5 | 28.54 | < 0.05 |
| | Depth × Temperature | 25 | 0.09 | 1.00 | 25 | 0.25 | 1.00 | 25 | 1.52 | 0.09 | 25 | 0.9 | 0.61 | 25 | 1.61 | 0.06 |
| LAP | Depth | 5 | 127.92 | < 0.05 | 5 | 64.27 | < 0.05 | 5 | 20.94 | < 0.05 | 5 | 77.8 | < 0.05 | 5 | 30.97 | < 0.05 |
|  | Temperature | 5 | 27.69 | < 0.05 | 5 | 29.61 | < 0.05 | 5 | 1.52 | 0.19 | 5 | 51.05 | < 0.05 | 5 | 69.4 | < 0.05 |
| | Depth × Temperature | 25 | 0.32 | 0.99 | 25 | 0.33 | 0.99 | 25 | 0.56 | 0.95 | 25 | 0.18 | 1.00 | 25 | 0.19 | 1.00 |
| AP | Depth | 5 | 44.43 | < 0.05 | 5 | 29.08 | < 0.05 | 5 | 25.22 | < 0.05 | 5 | 3.2 | 0.01 | 5 | 1.47 | 0.212 |
|  | Temperature | 5 | 9.72 | < 0.05 | 5 | 20.23 | < 0.05 | 5 | 1.53 | 0.19 | 5 | 7.61 | < 0.05 | 5 | 14.31 | < 0.05 |
| | Depth × Temperature | 25 | 0.15 | 1.00 | 25 | 0.26 | 1.00 | 25 | 0.32 | 0.99 | 25 | 0.09 | 1.00 | 25 | 0.15 | 1.00 |

**Figure 1.**
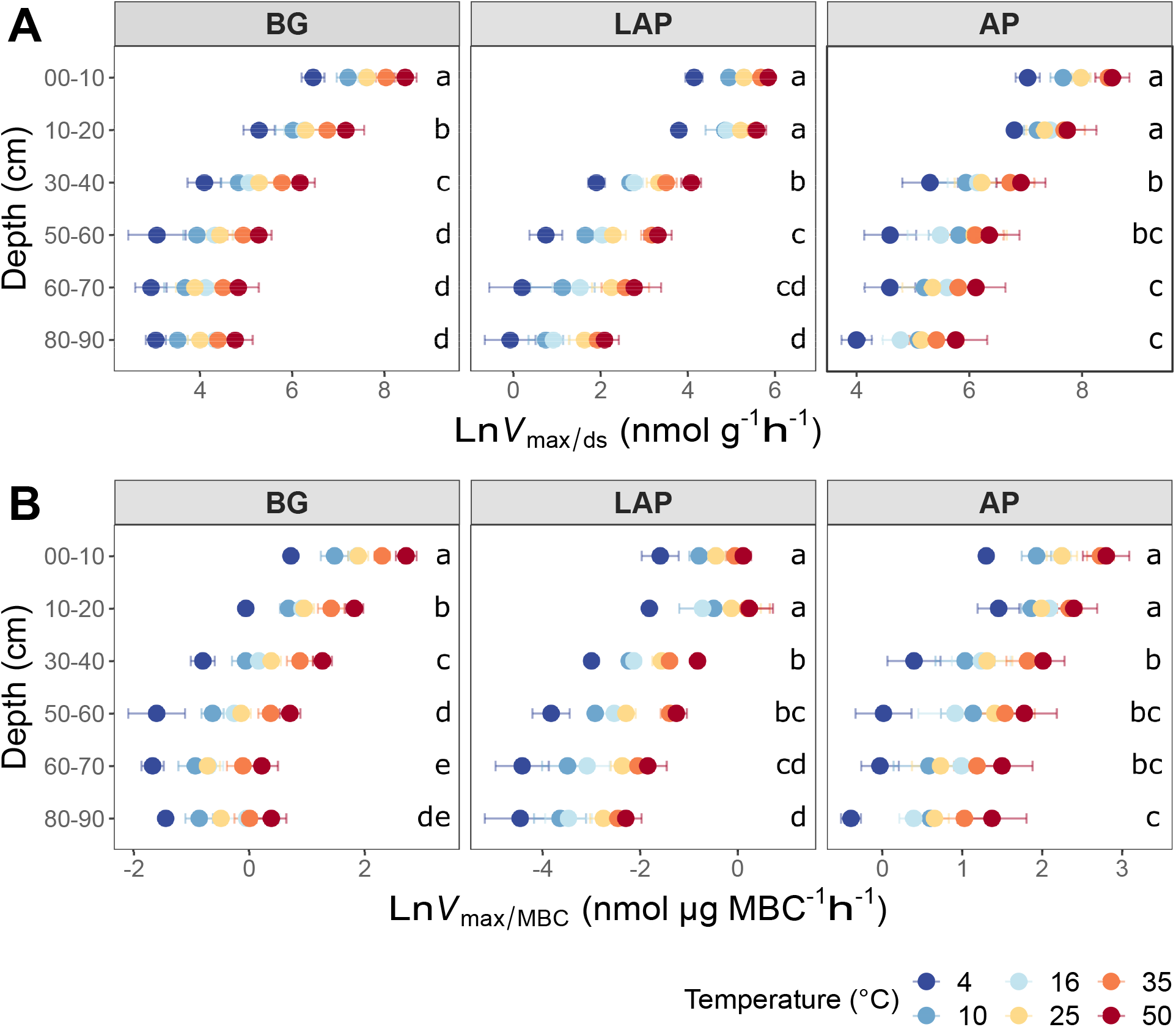
Enzyme maximum velocity (*V*_max_) at different depths and six temperatures from 4 to 50°C, expressed per (A) dry soil mass (*V*_max/ds_), or (B) microbial biomass C (*V*_max/MBC_). Colors indicate incubation temperatures and letters indicate significant differences (*p* < 0.05) between depths, per enzyme. Significance is based on Tukey’s tests after two-way ANOVA with depth and temperature as interactive factors (Table 2). Error bars represent the standard error of the mean (n = 3).

Since the concentration of microbial biomass carbon (MBC) declined strongly with soil depth, especially over the upper 30 cm (Figure S1), much of the decline in *V*_max/ds_ with depth may have been driven by lower microbial abundance. Therefore, we computed a biomass-specific *V*_max_, by expressing it per unit MBC (*V*_max/MBC_). *V*_max/MBC_ of all enzymes declined significantly down the soil profile over all temperatures, following the same trends as those of *V*_max/ds_ (*p* < 0.001) (Tables 2,S2, Figure 1B). The overall decline in *V*_max/MBC_ between 10-20 and 60-90 cm was only 8% lower (88.4 ± 1.7%) than that of *V*_max/ds_, indicating that variation in *V*_max/ds_ did not depend primarily on microbial biomass concentration. Like *V*_max/ds_, *V*_max/MBC_ did not show a significant interaction between depth and temperature. Also similar to *V*_max/ds_, *V*_max/MBC_ of AP and LAP did not vary within the upper 20 cm and declined mostly from 20 to 30 cm (Figure 1B). *V*_max/MBC_ did not decline significantly over the mid-depth range for either AP or LAP, but it was significantly lower at 80-90 cm than at 30-40 cm for AP, and lower between 60 and 90 cm than at 30-40 cm for LAP. *V*_max/MBC_ of BG declined more consistently down to 70 cm over all temperatures, but did not vary further.

*K*_m_ also declined (i.e., enzyme affinity increased) significantly with depth for all enzymes across temperatures (*p* < 0.001) with no significant interaction between depth and temperature (Tables 2,S2). However, *K*_m_ declined less with depth than *V*_max/ds_ or *V*_max/MBC_, and mainly between the upper 20 cm and lower depths, by up to 85.6 ± 1.3%, with some differences between enzymes. *K*_m_ of AP and BG declined with depth following trends similar to those of their *V*_max_ (Figure 2): *K*_m_ of AP declined mainly from 20 to 30 cm and remained relatively constant down to 80 cm, although it was significantly lower at 80-90 cm than at 30-40 cm; *K*_m_ of BG declined consistently down to 40 cm, without further variation (despite a spuriously high mean *K*_m_ at 80-90 cm only at 16°C). Unlike its *V*_max_, the *K*_m_ of LAP only declined from 20 to 30 cm, and did not vary significantly below that depth.

**Figure 2.**
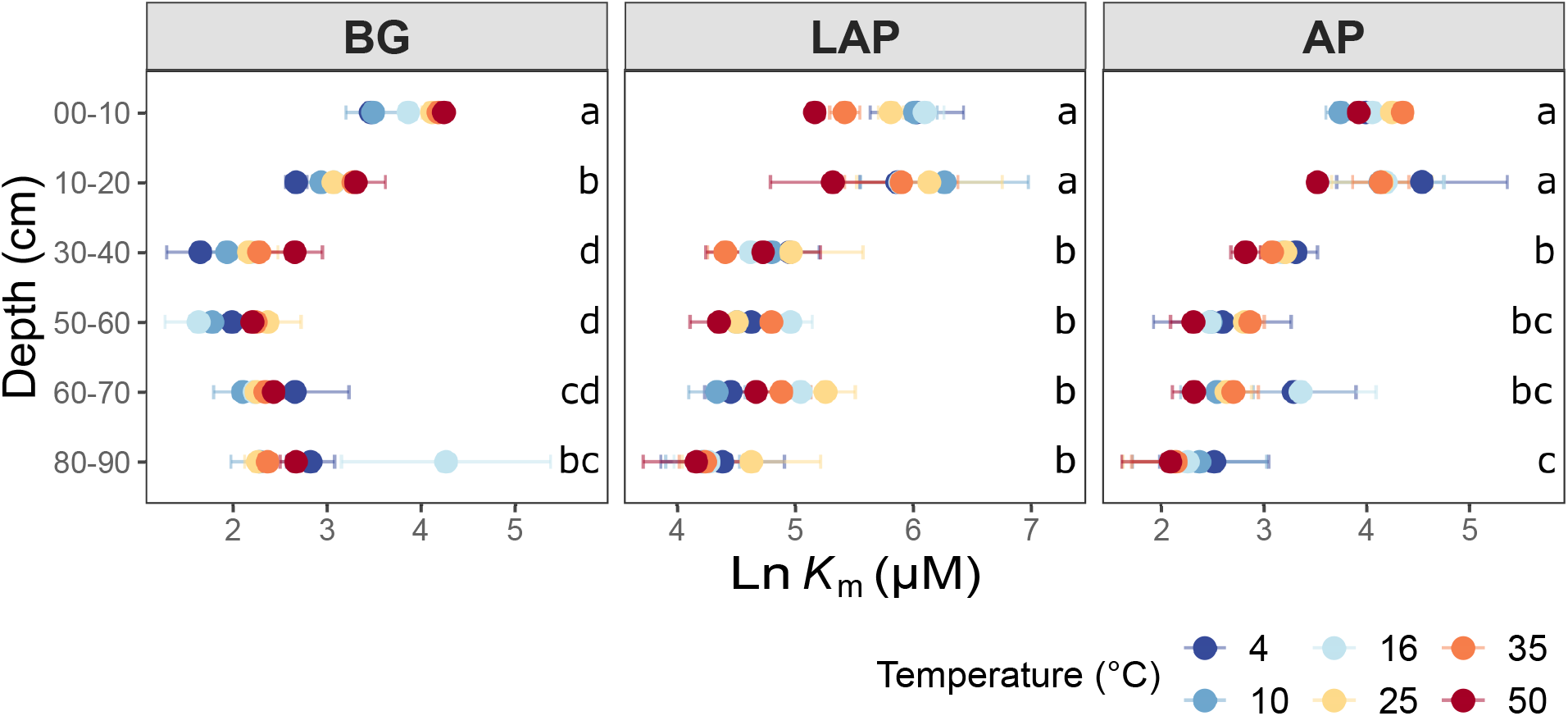
Enzyme Michaelis constant (*K*_m_) at different depths and six temperatures from 4 to 50°C. Colors indicate incubation temperatures and letters indicate significant differences (*p* < 0.05) between depths, per enzyme. Significance is based on Tukey’s tests after two-way ANOVA with depth and temperature as interactive factors (Table 2). Error bars represent the standard error of the mean (n = 3).

The apparent (i.e., observed) catalytic efficiency (CE_ds_), estimated as the ratio between *V*_max/ds_ and *K*_m_, reflects the catalytic efficiency of the enzyme pool present per mass of soil, regardless of microbial abundance or enzyme demand. CE_ds_ declined significantly across temperatures for all enzymes (*p* < 0.001), also without a significant interaction between depth and temperature (Tables 2,S2). The CE_ds_ of BG and LAP followed similar trends and declined mainly over the mid-depth range by up to 90.5 ± 1.1% over all temperatures, without significant variation in the upper 20 cm, or below 60 cm (Figure 3A). CE_ds_ of AP declined much less through the soil profile (58.1 ± 3.0%), and it was only significantly lower at 60-90 cm than in the upper 10 cm. Biomass-specific catalytic efficiency (CE_MBC_), estimated as the ratio between *V*_max/MBC_ and *K*_m_, represents the inherent catalytic efficiency of the enzyme pool produced by the local microbiome, as a function of its specific enzyme production capacity and demand. CE_MBC_ declined significantly with depth for BG and LAP (*p* < 0.001) across temperatures, but not for AP (Tables 2,S2, Figure 3B). CE_MBC_ of BG and LAP varied less with depth than other kinetic properties, and declined significant mainly below 60 cm by up to 71.8 ± 2.5% (Figure 3B). These declines in CE_MBC_ reflected the decline in *V*_max/MBC_ at lower depths, where *K*_m_ remained relatively constant. Like for all other kinetic parameters, depth-dependent differences in CE_MBC_ were not dependent on temperature (Table 2).

**Figure 3.**
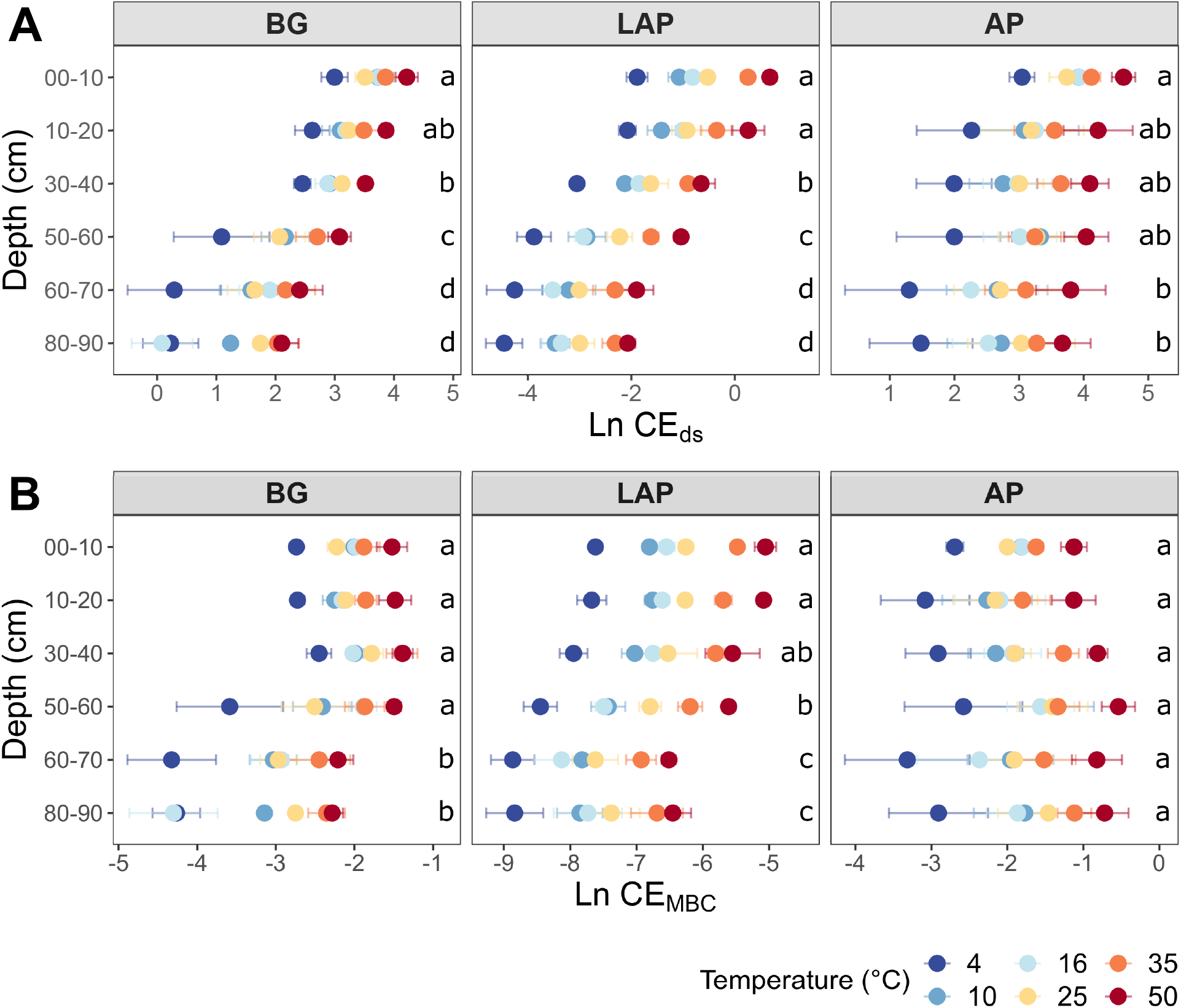
Enzyme catalytic efficiency at different depths and six temperatures from 4 to 50°C, calculated on a (A) dry soil mass basis (CE_ds_), or a (B) microbial biomass C basis (CE_MBC_). Colors indicate incubation temperatures and letters indicate significant differences (*p* < 0.05) between depths, per enzyme. Significance is based on Tukey’s tests after two-way ANOVA with depth and temperature as interactive factors (Table 2). Error bars represent the standard error of the mean (n = 3).

As there was no significant interaction between depth and temperature for any kinetic parameter (Table 2), we performed a random effects ANOVA with depth as independent variable and temperature as block variable, to control for possible confounding effects of the latter on depth-dependent differences. This analysis yielded the same results as the fixed effects ANOVA reported above, with the single exception that *V*_max/ds_ of AP declined significantly also between 50-60 cm and 80-90 cm (data not shown).

### Exoenzyme kinetics vary among enzymes as a function of soil depth

Variation in kinetic parameters among enzymes and interactions with temperature were tested using two-way fixed effects ANOVA per depth, with enzyme and temperature as independent factors, followed by post-hoc Tukey’s tests.

All kinetic properties varied significantly among enzymes at all depths and across temperatures (*p* < 0.001), but there was only a significant interaction between enzyme and temperature in the upper 10 cm for *K*_m_, CE_ds_ and CE_MBC_ (*p* < 0.001) (Table S3). *V*_max_ differed significantly among all enzymes at all depths, with AP having consistently the highest values, followed by BG and then LAP (*p* < 0.05) (Table S4, Figure 1A-B). *K*_m_ differed significantly among all enzymes between 10 and 60 cm (*p* < 0.05), but it did not differ between AP and BG in the upper 10 cm or below 60 cm (Table S4, Figure 2). LAP had always the highest *K*_m_ (i.e., lowest affinity). In contrast, BG had always the lowest *K*_m_, at least at depths where it was significantly different than that of AP (i.e., between 10 and 60 cm). CE_MBC_ differed significantly among all enzymes in the upper 10 cm and bellow 50 cm (*p* < 0.05), but not between AP and BG from 10 to 40 cm (Table S4, Figure 3B). Similar to *V*_max_, the CE_MBC_ of LAP was always the lowest among enzymes, followed by those of BG and then AP at depths where it differed significantly (Table S4, Figure 3B). As CE_MBC_ was calculated based on the same MBC value for all enzymes at each depth, CE_ds_ varied between enzymes similarly to CE_MBC_ (Table S4, Figure 3A).

We investigated the ratios between kinetic properties (*V*_max_, *K*_m_, and CE) of the three different enzymes as indicators for variation in relative nutrient demand through the soil profile. Based on two-way fixed effects ANOVA with depth and temperature as independent factors, all kinetic ratios between enzymes varied significantly with depth over all temperatures, with the exception of ratios between *K*_m_ of LAP and AP (*K*_m_^LAP:AP^) (*p* < 0.005) (Table S5). Temperature had a significant, positive effect on the *V*_max_ ratio of BG to AP (*V*_max_^BG:AP^), and LAP to AP (*V*_max_^LAP:AP^), and a significant, negative effect on ratios between catalytic efficiencies of BG and LAP (CE^BG:LAP^) (*p* < 0.05). Only CE^BG:LAP^ showed a depth × temperature interaction (*p* < 0.05) (Table S5, Figure 4A-I). *V*_max_^BG:LAP^ declined significantly from 10 to 20 cm, followed by a suggestive continuous increase down to 90 cm, although it was only significantly higher at 80-90 cm than at 10-20 cm (Figure 4A). *V*_max_^BG:AP^ declined with depth down to 70 cm, mainly from 10 to 20 cm and from 40 to 60 cm, followed by a significant increase from 70 to 90 cm that appeared to result partially from a spurious high value only at 16°C, among all six temperatures (Figure 4B). *V*_max_^LAP:AP^ generally declined with depth below 20 cm, but this variation was mainly significant between the upper 20 cm and the lower 30 cm (i.e., from 60 to 90 cm) (Figure 4C). *K*_m_^BG:LAP^ and *K*_m_^BG:AP^ followed trends similar to those of their corresponding *V*_max_ ratios: *K*_m_^BG:LAP^ declined significantly from 10 to 20 cm and remained relatively invariant through the profile (Figure 4D), whereas *K*_m_^BG:AP^ declined from 10 to 20 cm, followed by a suggestive but non-significant increase down to 70 cm (Figure 4E). The significant increases in *K*_m_^BG:LAP^ and *K*_m_^BG:AP^ at 80-90 cm were likely driven, at least partially, by the same spurious high *K*_m_ value of BG only at 16°C mentioned above. In contrast, *K*_m_^LAP:AP^ did not vary significantly with depth nor show any apparent trends (Figure 4F). CE^BG:LAP^ increased significantly from the upper 20 cm to a depth of 30-40 cm, although below that depth it did not vary significantly from any upper depths (Figure 4G). CE^BG:AP^ did not vary within the upper 40 cm, but it was significantly lower below 50 cm (Figure 4H). CE^LAP:AP^ declined with depth below 20 cm, but this variation was only significant between the upper 40 cm and the lower 30 cm (from 60 to 90 cm) (Figure 4I). This variation in CE^LAP:AP^ with depth mirrored the general trend of *V*_max_^LAP:AP^, as *K*_m_^LAP:AP^ was relatively invariant though the soil profile.

**Figure 4.**
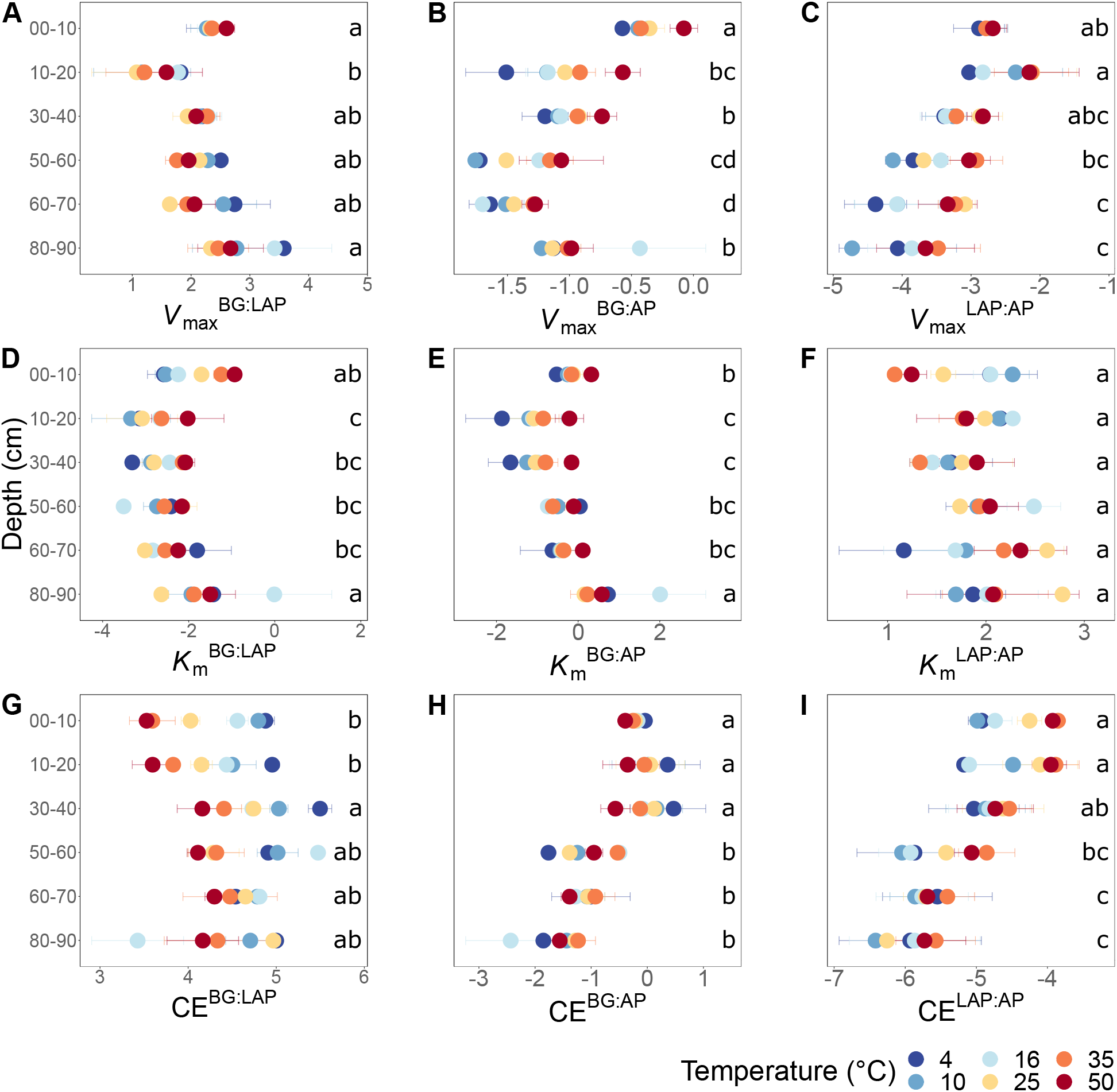
Ratios between kinetic parameters of BG, LAP and AP at different depths and six temperatures from 4 to 50°C. (A) *V*_max_^BG:LAP^, (B) *V*_max_^BG:AP^, (C) *V*_max_^LAP:AP^, (D) *K*_m_^BG:LAP^, (E) *K*_m_^BG:AP^, (F) *K*_m_^LAP:AP^, (G) CE^BG:LAP^, (H) CE^BG:AP^, and (I) CE^LAP:AP^. Colors indicate incubation temperatures and letters indicate significant differences (*p* < 0.05) between depths, per ratio. Significance is based on Tukey’s tests after two-way ANOVA with depth and temperature as interactive factors (Table S5). Error bars represent the standard error of the mean (n = 3); dots without error bars represent data-points with n < 3 (data excluded due to Michaelis-Menten models with poor fit).

### Comparison of temperature sensitivity based on Arrhenius and Macromolecular Rate Theory models

The temperature sensitivity of enzyme *V*_max_ was determined using a linear Arrhenius model over the full temperature range tested (4-50°C, *n* = 6) (Figure S2) and a realistic *in situ* soil range (4-35°C, *n* = 5) (Figure 5), as well as the non-linear MMRT model over the full temperature range (4-50°C, *n* = 6). The great majority of individual MMRT models (n = 54) approached the expected unimodal behavior, whose highest point represents *T*_opt_ (Figure 6). Despite the relatively high adjusted R^2^ values of the linear Arrhenius models over the full temperature range (0.81 ± 0.02, mean ± se) (Table S6), there was a general decline in the response rate of *V*_max_ between the two highest temperatures consistent with MMRT predictions, even when *T*_opt_ was not reached (Figure 6). The temperature response of *V*_max_ in four of the 54 individual models did not fit the assumptions of MMRT, resulting in biologically implausible *T*_opt_ and TS_max_ estimates for these soils, namely negative values due to an upward concave response (AP at 10-20 cm and BG at 50-60 cm, replicate 3), or values above 200°C due to approximately linear responses (BG at 60-70 cm, replicate 1; and BG at 10-20 cm, replicate 3) (Figure 6, Table S7). These model estimates were thus excluded from further analyses. Comparisons between alternative Arrhenius and MMRT models based on the Akaike Information Criterion (AIC), corrected AIC (AICc) and Bayesian Information Criterion (BIC) standard indices indicated that, in general, their model fit did not differ substantially (see also Supplementary Results). Nevertheless, overall model comparisons tended to favor MMRT, which further suggested that MMRT provided a more realistic representation of the temperature response of enzyme *V*_max_.

**Figure 5.**
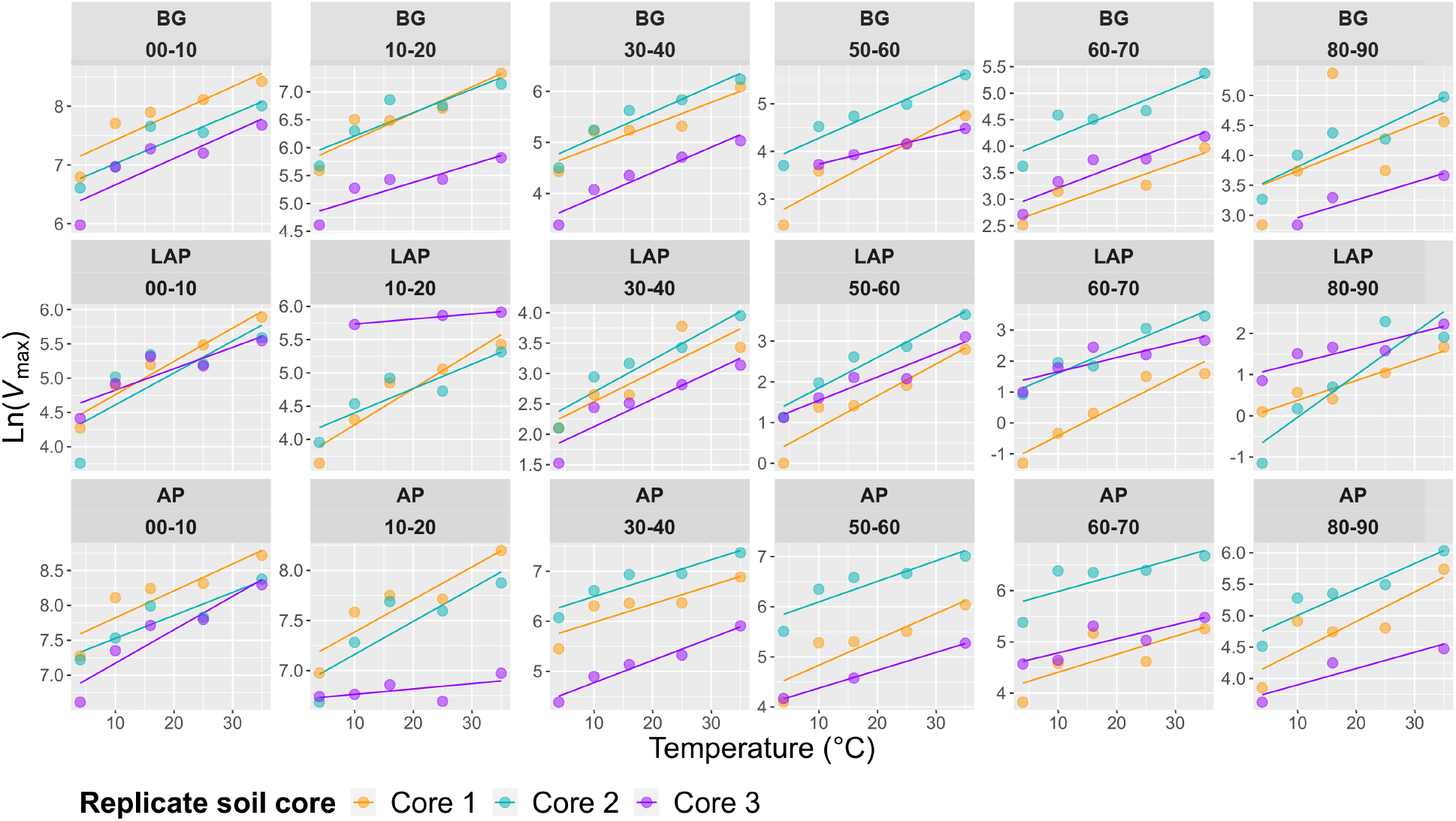
Arrhenius models of *V*_max_ over five temperatures from 4 to 35°C per enzyme, depth and replicate core sample.

**Figure 6.**
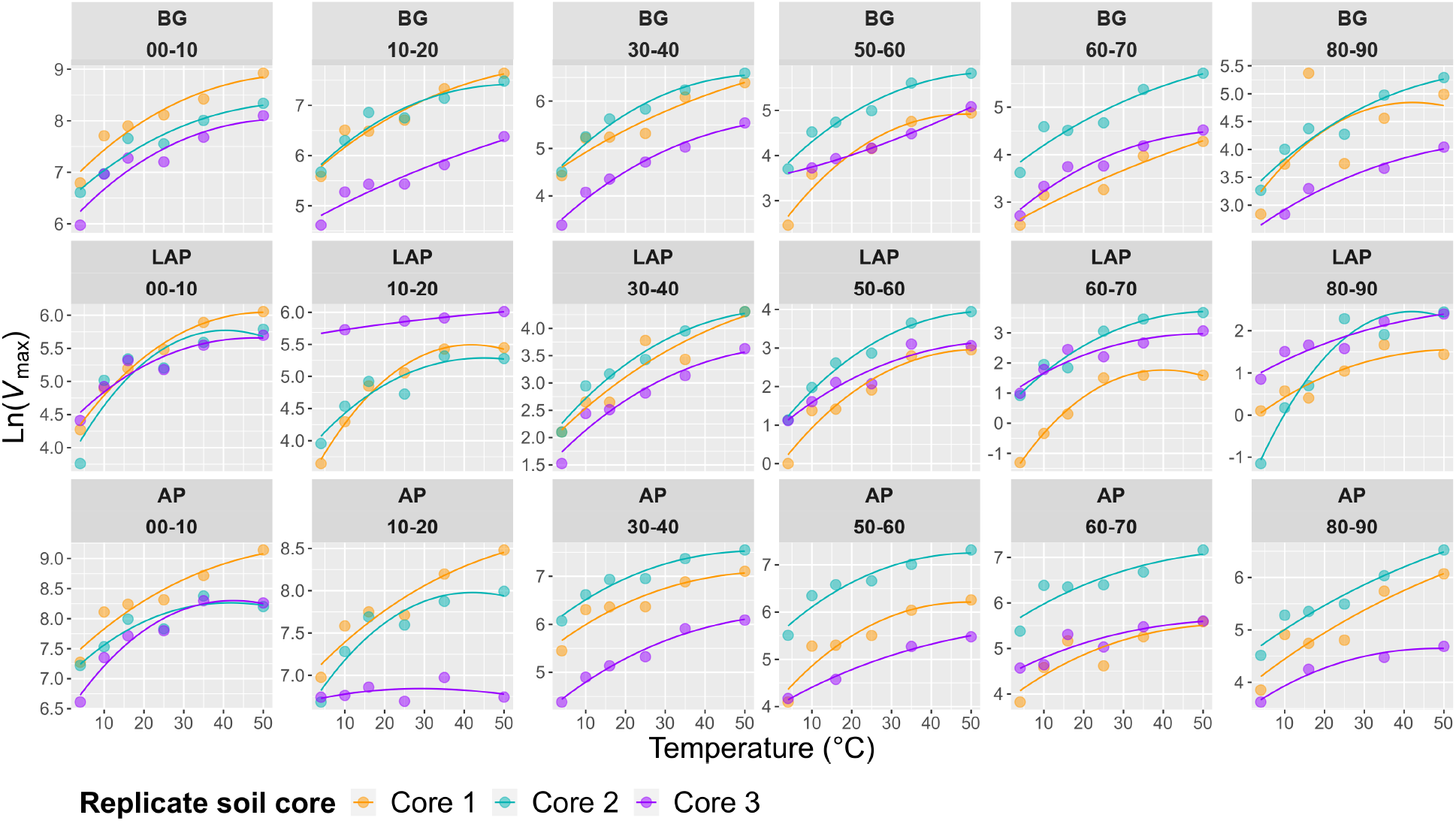
MMRT models of *V*_max_ over six temperatures from 4 to 50°C per enzyme, depth and replicate core sample.

### Temperature sensitivity of exoenzymes is similar through the soil profile

Temperature had an overall significant effect on *V*_max_, CE_MBC_ and CE_ds_ of all enzymes, and on *K*_m_ of only BG, over the whole soil profile (*p* < 0.001) (Table 2, Figures 1-3). Enzyme temperature sensitivity was further assessed based on Q_10_ coefficients of *V*_max_ (*V*_max/MBC_ and *V*_max/ds_ scale identically with temperature at each depth), *K*_m_ and CE (CE_MBC_ and CE_ds_ scale identically with temperature at each depth) following the Arrhenius model assumption of linearity, and on the temperature optimum (*T*_opt_), point of maximum temperature sensitivity (TS_max_) and change in heat capacity (Δ*C*_p_^‡^) estimated using the non-linear MMRT model. Differences in temperature sensitivity parameters between depths or enzymes were tested with one-way ANOVA per enzyme or depth, respectively.

Despite some suggestive, depth-dependent trends in the Q_10_ values of *V*_max_, *K*_m_ and CE, they did not vary significantly with depth for any enzyme (Table 3), whether estimated over a realistic *in situ* soil temperature range (4-35°C) (Figure 7), or the full experimental temperature range (4-50°C) (Figure S3). However, Q_10_ values varied significantly between some enzymes in a depth-dependent manner (Tables S9-S10). Both temperature ranges yielded similar Q_10_ values that varied within a narrow range, although Q_10_ values over 4-50°C were slightly lower than those over 4-35°C due to a frequent decline in the response rate of *V*_max_ between 35 and 50°C (Table S8). Therefore, only Q_10_ values over the realistic *in situ* soil temperature range are henceforth presented. The Q_10_ of *V*_max_ was consistently above 1, indicating a positive effect of temperature on *V*_max_ across depths of 1.44 ± 0.03, 1.56 ± 0.03 and 1.78 ± 0.10 (mean ± se) for AP, BG and LAP, respectively (Figure 7A, Table S8). Mean activation energies (*E*_a_) estimated from the same linear relationships were 25.69 ± 1.74, 31.50 ± 1.45, and 39.33 ± 3.98 kJ mol^-1^ K^-1^ (mean ± se) across depths for AP, BG and LAP respectively (Table S8). Between enzymes, the Q_10_ of *V*_max_ was only significantly different between AP and LAP at 60-70 cm (*p* < 0.05) (Table S10). *K*_m_ was relatively insensitive to temperature, with mean Q_10_ values across depths of 1.00 ± 0.06, 1.14 ± 0.05 and 0.99 ± 0.07 (mean ± se) for AP, BG and LAP, respectively (Figure 7B, Table S8). This was consistent with the two-way ANOVA showing that temperature had generally no significant effect on *K*_m_. The significant effect of temperature on the *K*_m_ of BG detected by the two-way ANOVA was likely due to the spurious high *K*_m_ of BG only at 16°C at 80-90 cm (Table 2, Figure 2), which was not reflected on the Q_10_ computed across temperatures. Although the mean Q_10_ of *K*_m_ was similar between enzymes and depths, it was significantly lower for LAP in the upper 10 cm relative to the other two enzymes (*p* < 0.05) (Figure 7B, Table S10). This difference suggested that the affinity of LAP might increase with warmer temperatures (i.e., lower *K*_m_) at this depth (Q_10_ = 0.83 ± 0.07, mean ± se), compared to those of AP (Q_10_ = 1.18 ± 0.06) or BG (Q_10_ = 1.30 ± 0.07). The Q_10_ of CE was consistently above 1, similar to that of *V*_max_, with similar overall mean values across enzymes: 1.51 ± 0.08, 1.42 ± 0.08 and 1.82 ± 0.05 (mean ± se) for AP, BG and LAP respectively (Figure 7C, Table S8). However, the Q_10_ of CE was significantly higher for LAP in the upper 10 cm (1.86 ± 0.07, mean ± se) relative to those of AP (1.27 ± 0.02) or BG (1.20 ± 0.08) (*p* < 0.05) (Figure 7C, Table S10), reflecting the apparent negative effect of higher temperatures on the *K*_m_ of LAP at that depth.

**Table 3.**
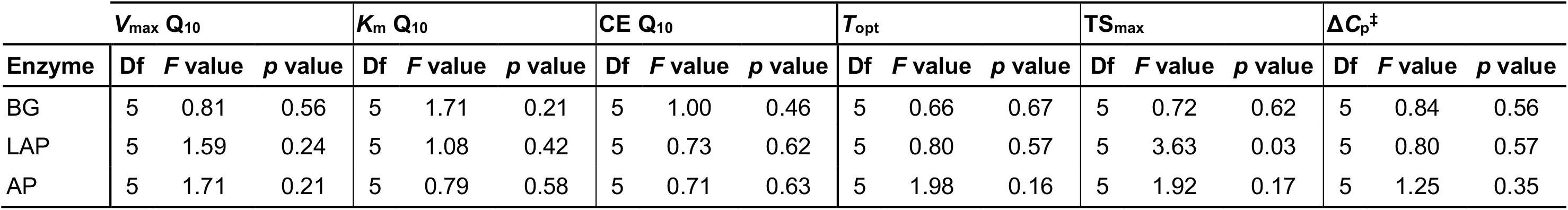
One-way ANOVA of temperature sensitivity estimates with depth. Q_10_ values were calculated between 4-35°C, and the MMRT model parameters *T*_opt_, TS_max_, and Δ*C*_p_^‡^ between 4-50°C. Differences were considered significant at *p* < 0.05.

| | $V_{max} Q_{10}$ | | | $K_m Q_{10}$ | | | $CE Q_{10}$ | | | $T_{opt}$ | | | $TS_{max}$ | | | $\Delta C_p^\ddagger$ | | |
| --- | --- | --- | --- | --- | --- | --- | --- | --- | --- | --- | --- | --- | --- | --- | --- | --- | --- | --- |
| Enzyme | Df | F value | p value | Df | F value | p value | Df | F value | p value | Df | F value | p value | Df | F value | p value | Df | F value | p value |
| BG | 5 | 0.81 | 0.56 | 5 | 1.71 | 0.21 | 5 | 1.00 | 0.46 | 5 | 0.66 | 0.67 | 5 | 0.72 | 0.62 | 5 | 0.84 | 0.56 |
| LAP | 5 | 1.59 | 0.24 | 5 | 1.08 | 0.42 | 5 | 0.73 | 0.62 | 5 | 0.80 | 0.57 | 5 | 3.63 | 0.03 | 5 | 0.80 | 0.57 |
| AP | 5 | 1.71 | 0.21 | 5 | 0.79 | 0.58 | 5 | 0.71 | 0.63 | 5 | 1.98 | 0.16 | 5 | 1.92 | 0.17 | 5 | 1.25 | 0.35 |

**Figure 7.**
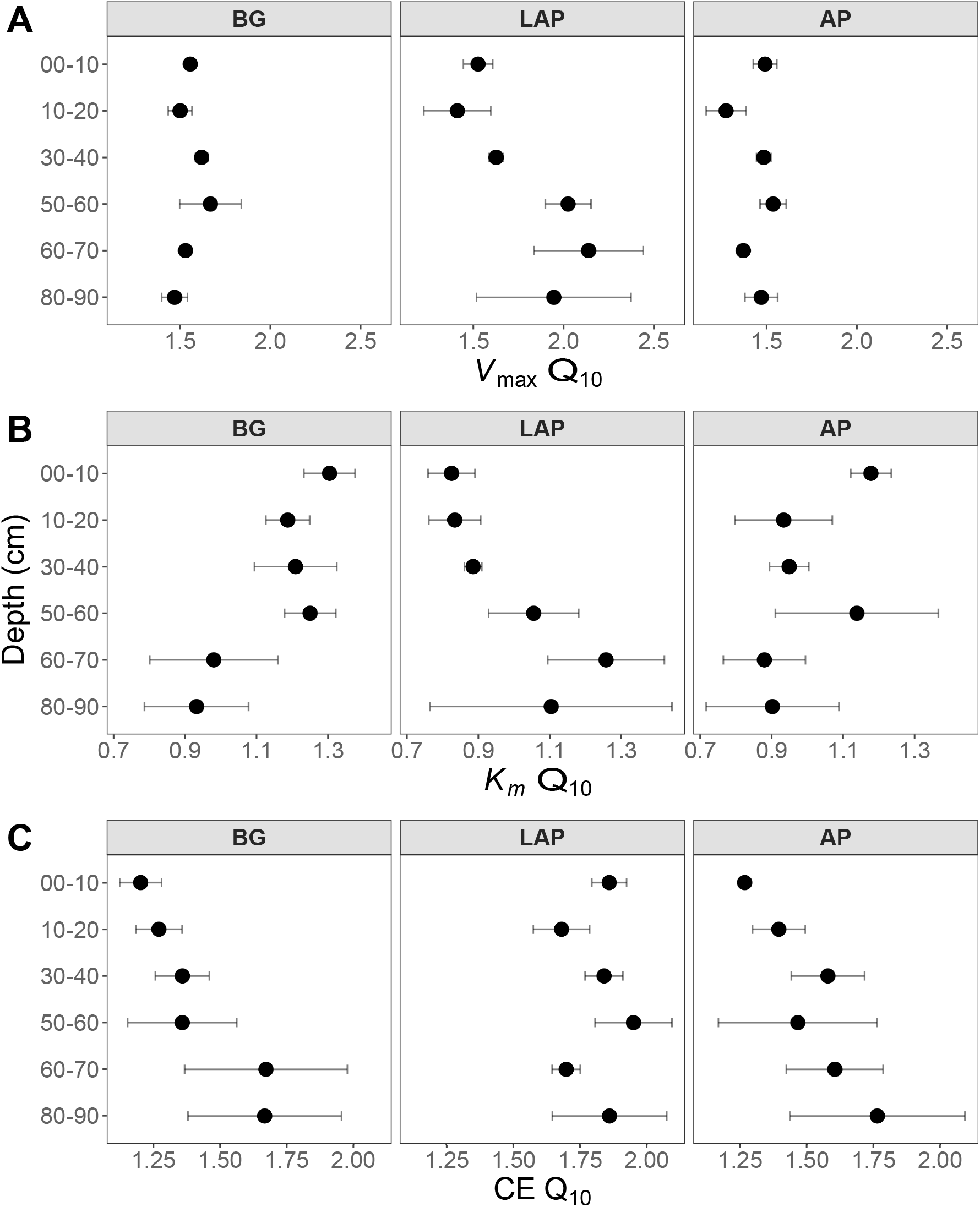
Q_10_ of Michaelis-Menten kinetic parameters over five temperatures from 4 to 35°C at different depths. (A) *V*_max_, (B) *K*_m_, and (C) CE. Differences between depths are not significant (*p* > 0.05), based on ANOVA per enzyme (Table 3). Error bars represent the standard error of the mean (n = 3).

None of the temperature sensitivity parameters estimated by MMRT –*T*_opt_, TS_max_ and Δ*C*_p_^‡^– varied significantly with depth (with one exception; see below), or between enzymes (Figure 8, Tables 3,S9). These parameter estimates showed considerable variability among replicates, and estimates from models with poor fit to MMRT’s predicted behavior (*T*_opt_ or TS_max_ < 0°C, or > 200°C; four out of 54 total models) were excluded from the analysis, likely reducing the statistical power of few pairwise comparisons between depths and enzymes. Mean *T*_opt_ and TS_max_ were consistent across depths and enzymes, with mean values of 65.19 ± 3.74°C and 31.63 ± 1.98°C (mean ± se), respectively (Figure 8A-B, Table S8). TS_max_ of LAP was significantly different between the 0-10 and 30-40 cm depth intervals (*p* < 0.05), which was the only exception to otherwise non-significantly different parameter estimates across either enzymes or depths. Mean Δ*C*_p_^‡^ values were similar between AP and BG across depths, with a combined mean value of -0.79 ± 0.06 kJ mol^-1^ K^-1^ (mean ± se) (Figure 8C, Table S8). Δ*C*_p_^‡^ of LAP spanned a broader range of values (−1.32 ± 0.21, mean ± se), mainly due to suggestive, albeit non-significant, lower values at depths below 60 cm.

**Figure 8.**
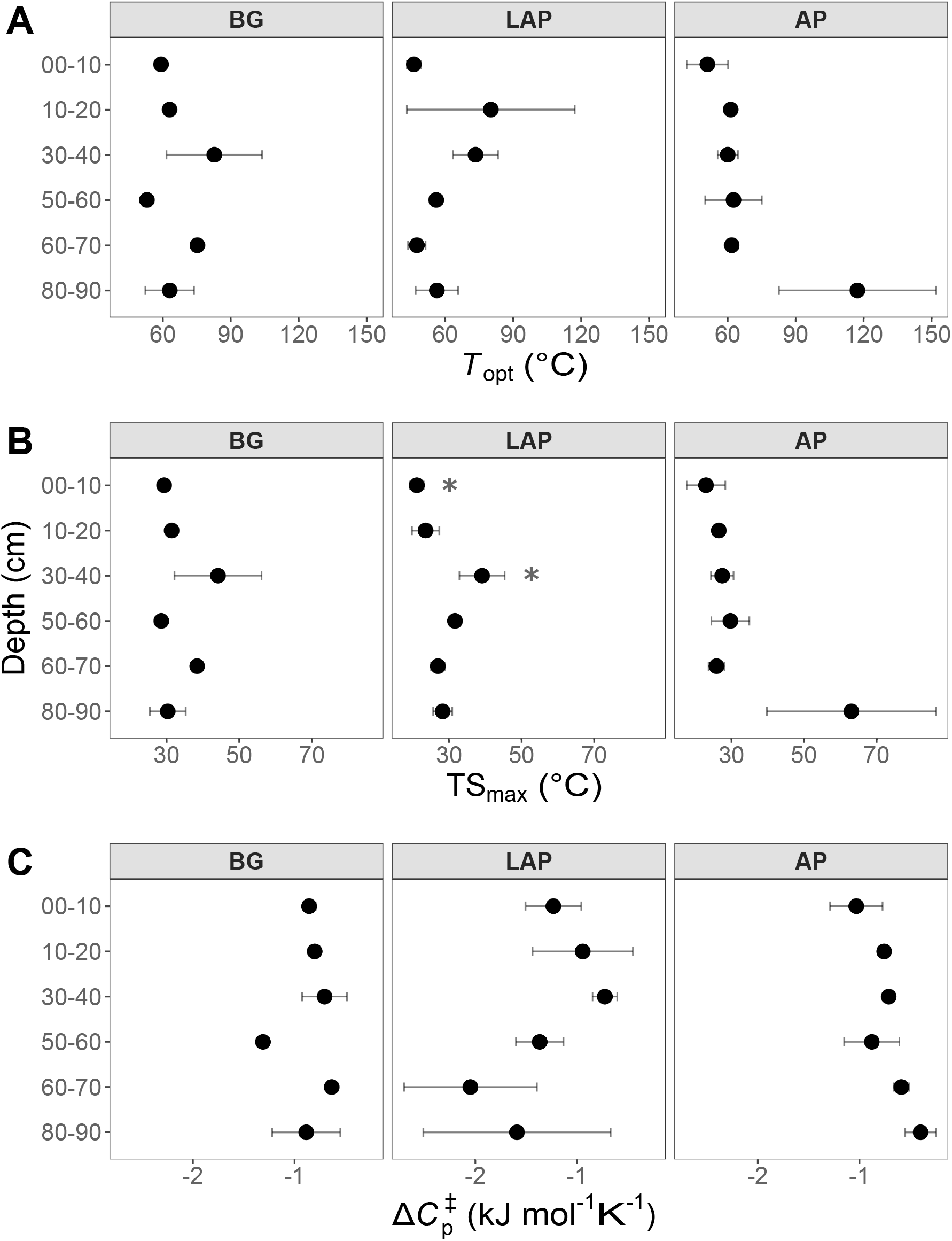
MMRT model estimates over six temperatures from 4 to 50°C at different depths. (A) *T*_opt_, (B) TS_max_, and (C) Δ*C*_p_^‡^. Differences between depths are not significant (*p* > 0.05), based on ANOVA per enzyme (Table 3), except for TS_max_ of LAP between the depths indicated by asterisks (*). Error bars represent the standard error of the mean (n = 3).

## Discussion

We show that kinetic properties of the enzymes BG, LAP and AP varied markedly through the soil profile at a temperate forest, even when accounting for the large variation in microbial biomass. Moreover, this variation in enzyme kinetics was independent from temperature, as kinetic properties varied similarly between soil depths over temperatures between 4 and 50°C. We also show that the temperature sensitivity of these enzymes is similar through the soil profile, based on both linear Arrhenius and non-linear MMRT models, although temperature can directly affect the relative kinetics between enzyme types. To our knowledge, this is the first study to investigate the MM kinetic properties of soil enzymes and their direct temperature sensitivity through the soil profile, in this case to 90 cm.

### Higher enzyme affinity, but lower *V*_max_ and catalytic efficiency, indicate adaptation to lower substrate availability and distinct microbial life strategies in deeper soils

As hypothesized, *V*_max/ds_ and *K*_m_ declined strongly with soil depth, but followed distinct trends that depended on enzyme type. The decline in *V*_max/ds_ of 96.4 ± 0.4% across enzymes and temperatures down to 90 cm indicated a drastic decline in enzyme production capacity, product demand, or substrate availability. This was consistent with the lower microbial biomass concentrations in deeper soils observed here, as generally reported across studies (Blume et al., 2002; Fierer et al., 2003a; Jones et al., 2018; Loeppmann et al., 2016a; Schnecker et al., 2014). Higher density of plant roots in surface soils may have also contributed to higher near-surface *V*_max/ds_, as shown particularly for BG in rooted soils when compared to fallow soils (Loeppmann et al., 2016b), and in rhizosphere hotspots when compared to bulk soil (Tian et al., 2020). Plants may induce higher enzyme *V*_max_ by promoting microbial growth through C-rich exudates, competing with microbes for N and P, or directly stimulating microbial enzyme production to enhance availability of assimilable products (Burns et al., 2013). Declines in activity of hydrolytic enzymes have been consistently observed down to depths between 50 and 420 cm in diverse soils, including from temperate, taiga and tropical forests, arctic tundra, grasslands and croplands (Darby et al., 2020; Dove et al., 2020; Gelsomino and Azzellino, 2011; Jing et al., 2017; Kramer et al., 2013; Loeppmann et al., 2016a; Schnecker et al., 2014, 2015; Stone et al., 2014; Taylor et al., 2002; Venkatesan and Senthurpandian, 2006). However, among the five studies from which we could retrieve at least approximate *V*_max/ds_ values, only Loeppmann et al. (2016a) observed mean declines in *V*_max/ds_ of BG, LAP and AP down to 70 cm similar to those observed here down to 90 cm (91.0 ± 2.2%), whereas others observed substantially smaller mean declines of approximately 67.5 ± 6.3% for AP and BG at depths between 55 and 110 cm (Gelsomino and Azzellino, 2011; Stone et al., 2014; Venkatesan and Senthurpandian, 2006). Consistent with this, only Loeppmann et al. (2016a) determined *V*_max_ based on a MM model over a series of enzyme substrate concentrations, as needed to estimate MM kinetics accurately. Potential enzyme activities have been frequently shown to correlate positively with MBC concentration (Gelsomino and Azzellino, 2011; Perucci, 1992; Ren et al., 2018; Stone et al., 2014), as *V*_max_ is linearly dependent on enzyme concentration, which in turn is largely dependent on microbial abundance. However, this is not always the case (Waring et al., 2014), as exoenzyme production may also be induced or repressed depending on substrate availability and product demand, following the evolutionary-economic mechanisms that regulate allocation of cellular resources (Allison et al., 2011; German et al., 2011a). Moreover, enzyme production varies between organisms and is subject to variable levels of regulation (Allison et al., 2011; Burns et al., 2013; Sinsabaugh and Shah, 2012). For example, some isozymes and enzyme types may be expressed at stable constitutional levels under specific conditions, as previously suggested for deep soils (Stone et al., 2014), whereas others may be more strictly responsive to cellular demands or environmental cues. Therefore, a biomass-specific *V*_max_ (*V*_max/MBC_) can be interpreted as a catalytic rate constant independent of microbial abundance, which represents the collective effect of inherent enzyme properties, and specific enzyme production and demand of the microbiome. Surprisingly, *V*_max/MBC_ of all enzymes declined nearly as much with depth as *V*_max/ds_, indicating that microbial abundance was not the primary driver of variation in *V*_max_. Loeppmann et al. (2016a) found similar trends only for BG and LAP, and only below 30-40 cm, as *V*_max/MBC_ increased from the surface to that depth and only then declined consistently down to 70 cm. These differences might have resulted from a steeper decline in substrate availability and microbial biomass in our soils, which are covered by a thick litter layer and have a shallow rhizosphere, compared to a more extensive rhizosphere (i.e., maize) and less surface litter in the soils studied by Loeppmann et al. (2016a). The consistent decline in *V*_max/MBC_ in deeper soils observed by us, and to some extent also Loeppmann et al. (2016a), contrasts with most other studies where *V*_max/MBC_ of AP, BG and LAP either increased, or did not vary with depth (Dove et al., 2020; Gelsomino and Azzellino, 2011; Kramer et al., 2013; Schnecker et al., 2015; Stone et al., 2014). We could only identify one study that detected a decline in *V*_max/MBC_ of BG (Taylor et al., 2002), although declines with depth have been more frequently observed for other enzymes (Gelsomino and Azzellino, 2011; Schnecker et al., 2015; Taylor et al., 2002). However, *V*_max_ in those studies was inferred from a single concentration of substrate across depths that was generally below the saturation point necessary to reach the *V*_max_ values we observed, particularly in surface soils. We suggest that the apparent increase, or lack of variation, in *V*_max/MBC_ previously observed may have resulted from underestimating *V*_max_ in surface soils. Our results indicate that exoenzyme pools in deeper soils have inherently low activity potential due to lower expression levels and/or lower catalytic capabilities than those in surface soils, possibly reflecting differences in microbial life strategies and substrate preferences.

The consistent increase in affinity (i.e., decrease in *K*_m_) of all enzymes with depth by 85.6 ± 1.3%, mainly between 20 and 60 cm, indicated a major decline in substrate availability at mid-depths consistent with the decline in DOC, TDN (Figures S4A-B) and total C and N in these soils (Hicks Pries et al., 2017, 2018). This supported our hypothesis that persistently low substrate concentrations in deep soils select for microbes encoding isozymes with lower *K*_m_ in order to maintain relatively constant maximal catalytic rates (Sinsabaugh et al., 2014). *K*_m_ values largely above physiologic substrate concentrations would render enzyme activity entirely dependent on substrate availability, which, under deep soil conditions, would lead to suboptimal rates and provide limited return to the investment on enzymes. Conversely, higher substrate availability in surface soils through plant litter inputs likely alleviates the selective pressure on enzymes with high affinity. Inputs of readily assimilable compounds through root exudation may further alleviate this pressure by reducing the relative importance of continuously maintaining maximal depolymerization rates (Allison et al., 2011), similar to what has been observed for substrate induced respiration (Blagodatskaya et al., 2009). The variation in enzyme affinities with depth observed here indeed appeared to reflect the overall decline in root density –both fine and coarse roots– below 40 cm at this site (Hicks Pries et al., 2018). The *K*_m_ of BG particularly mirrored the continuous steep decline in fine root density down to 40 cm (Hicks Pries et al., 2018), whereas those of AP and LAP did not vary within the upper 20 cm. This suggests that availability of easily metabolizable C-containing compounds exuded by fine roots may have a particular regulatory effect on depolymerization of cellulose (Allison et al., 2011; Allison and Vitousek, 2005) through selection of microbes encoding BG isozymes with distinct affinities. This hypothesis is further supported by the higher affinities of cellulose-degradation enzymes, including BG, observed in fallow soils relative to rooted soils (Loeppmann et al., 2016b), and in bulk soils relative to rhizosphere hotspots (Tian et al., 2020). Moreover, microbes producing BG in the rhizosphere have been shown to be distinct from those in the detritusphere (Nuccio et al., 2020).

The catalytic efficiency (CE), also referred to as specificity constant (Gelsomino and Azzellino, 2011), can be determined as *K*_cat_/*K*_m_ when *K*_m_ exceeds the concentration of substrate present, which is typically the case under physiological conditions (Berg et al., 2002; Koshland, 2002). Therefore, CE not only represents a fundamental functional property under direct selective pressure, but also evolutionary tradeoffs between maximum substrate turnover (*K*_cat_) and affinity, which are themselves subject to selection (Sinsabaugh et al., 2014). However, *K*_cat_ expresses the maximum amount of substrate converted per unit of time, per enzyme unit, and thus it cannot be directly inferred from enzyme assays in complex environmental samples, such as soils, where specific enzyme concentrations are generally unknown and hard to quantify. In these cases, an apparent CE (CE_ds_) has been estimated as *V*_max/ds_/*K*_m_ (Kujur and Kumar Patel, 2014; Loeppmann et al., 2016b, 2016a; Moscatelli et al., 2012; Razavi et al., 2016; Triebwasser-Freese et al., 2015), which represents the observed CE of the enzyme pool present per mass of soil, regardless of the specific production capacity and demand of the microbiome. In our soils, CE_ds_ of all enzymes declined significantly with depth, with CE_ds_ of BG and LAP declining consistently over the mid-depth (20 to 60 cm) by up to ∼90%, whereas that of AP varied much less. These results are consistent with those from the only other study that, to our knowledge, determined CE_ds_ in deep soils, where CE_ds_ of these same enzymes declined by 2-to 20-fold between the upper 40 cm and depths down to 70 cm (Loeppmann et al., 2016a). Tian et al. (2020) have also shown that CE_ds_ of BG and AP was higher in fertile soils than in nutrient-poor soils, consistent with higher CE_ds_ in surface soils with greater nutrient availability than deep soils. However, contrary to our initial expectations, CE based on biomass-specific *V*_max_ (CE_MBC_) either declined by up to ∼70% (BG and LAP) or did not vary (AP) with depth. We initially hypothesized that CE would increase with depth to maximize return on the investment in enzymes, given the scarce substrates provided by lower plant litter inputs and lower compensation by root exudates. Microbial communities adapted to these conditions would be expected to encode isozymes with higher affinity (i.e., lower *K*_m_) and/or produce more exoenzymes per unit biomass (i.e., higher *V*_max/MBC_) in a proportion that favors higher *V*_max/MBC_/*K*_m_ ratios (i.e., CE_MBC_). As the *K*_m_ values of BG and LAP were relatively invariable between 30 and 90 cm, their lower CE_MBC_ at depths below 60 cm was mainly driven by a decline in *V*_max/MBC_, suggesting that it was constrained by lower production of enzymes per unit biomass below that depth, rather than higher *K*_m_. On the other hand, enzymes may optimize *K*_cat_ in adaptation to environmental pressures (e.g., temperature) at the expense of *K*_m_, leading to conformational adaptations that reduce active site binding, which result in higher *K*_m_ and suboptimal catalytic efficiencies (Struvay and Feller, 2012). Therefore, a lower CE_MBC_ in deeper soils driven by lower *V*_max/MBC_ may also reflect enzymes with lower *K*_cat_, as a result of trade-offs with *K*_m_, rather than lower enzyme production. The contrasting lack of variation in CE_MBC_ of AP with depth, regardless of individual variation in *V*_max/MBC_ and *K*_m_, might have resulted from different factors and interactions related to variation in relative P availability and demand, and in regulation of AP expression. Alternatively, P may be primarily acquired from minerals rather than organic compounds (Alori et al., 2017), and thus the CE_MBC_ of AP alone does not directly reflect P demand.

Our results suggest that microbial communities in deep subsoils encode exoenzymes with intrinsically lower *K*_cat_, and/or produce less enzymes per cell than those in surface soils, leading to a lower emergent CE_MBC_. The expectation that *V*_max/MBC_ and CE_MBC_ would increase with depth assumes that microbiomes have largely redundant metabolic and elemental demands, and thus exoenzymes are optimized to provide nutrients in proportion to the size and demands of the community, as a function of nutrient availability (Allison et al., 2011). However, several studies have shown that microbiomes change markedly with soil depth (Eilers et al., 2012; Fierer et al., 2003b; Hansel et al., 2008; Hartmann et al., 2009; Jiao et al., 2018; Liu et al., 2019), with deep soils harboring less diverse and functionally distinct organisms (Brewer et al., 2019; Diamond et al., 2019; Dove et al., 2021; Yan et al., 2019). Life strategies that prioritize cellular maintenance over fast growth and maximal resource exploitation, and properties such as utilization of alternative substrates, storage compound production, and ability to sporulate or undergo dormancy may all affect exoenzyme properties, and possibly contribute to relax selective pressures on their kinetics (Ho et al., 2017; Ramin and Allison, 2019). Dove et al. (2021) have shown that deep soil microbes at our site have lower growth rates and lower carbon use efficiency than those at the surface, reflecting a lower nutrient demand and greater investment on cellular maintenance that may underlie the lower *V*_max/MBC_ and CE_MBC_ we observed. Moreover, the declining substrate availability with depth is expected to decrease the return on exoenzyme investment and favor alternative metabolic strategies that do not rely primarily on depolymerization of complex organic matter. Dove et al. (2021) have indeed shown that microbiomes in these deep soils have lower potential to degrade complex carbohydrates, similar to those in other soils (Diamond et al., 2019). Conversely, it has been shown that deep soil communities are enriched in organisms that can metabolize C1 and other low molecular weight C compounds, and use inorganic N forms as energy sources (Brewer et al., 2019; Diamond et al., 2019), as well as in taxa that comprise mainly chemoautotrophs (Brewer et al., 2019; Cao et al., 2012; Diamond et al., 2019; Eilers et al., 2012; Turner et al., 2017). Therefore, a smaller fraction of organisms relying on exoenzymes in deep soils is also likely to contribute to a lower emergent CE_MBC_ due to both lower overall biomass-specific enzyme production and lower competition for enzyme products within the community.

### Kinetics varied between enzymes and indicated an increasing demand in P with depth relative to C and N

As expected, *V*_max_ differed significantly between all three enzymes at every depth, reflecting fundamental differences in relative nutrient demand, as well as possible differences in enzyme properties and regulation. According to ecoenzymatic stoichiometry, ratios between *V*_max_ of hydrolytic enzymes involved in acquisition of C, N or P reflect the relative demand in these elements in relation to their availability, and thus the equilibrium between microbial biomass and SOM stoichiometry (Sinsabaugh and Shah, 2012). Here, both *V*_max_ and CE ratios between C-and N-acquiring enzymes (*V*_max_^BG:LAP^ and CE^BG:LAP^, respectively) were relatively constant through the soil profile, despite suggestive increases at lower depths, which were consistent with the trend in soil C:N ratio (Hicks Pries et al., 2018) and ratios between dissolved C and N pools (Figure S4C). Consistently, laboratory incubations with substrate amendments have also shown that the proximate C limitation of soil heterotrophic respiration is similar between shallow and deep soils at this site (Dove et al., 2021). In turn, *V*_max_ and CE ratios between BG:AP and LAP:AP decreased with depth, suggesting an increasing demand for P relative to either C or N, similar to previous observations of ratios between activities of C- and P-acquiring enzymes in both temperate and tropical soils (Loeppmann et al., 2016a; Stone et al., 2014). While Dove et al. (2021) have shown that the proximate nutrient limitation of respiration was generally higher in surface soils, they applied combined N and P amendments, which might have masked the effects of individual nutrients on respiration at different soil depths. Together, these observations contrasted with the generally expected increase in C limitation of heterotrophic growth with depth, following the typical declines in C:N and C:P ratios (Fierer et al., 2003a; Mooshammer et al., 2014; Ni et al., 2020; Taylor et al., 2002). Alternatively, it has been shown that microbes can use phosphorylated compounds primarily as a C source (Heuck et al., 2015), and thus higher *V*_max_ and CE of AP may rather indicate higher C demand in the absence of more favorable C sources in deep soils, as previous suggested (Stone et al., 2014). It should be noted, however, that depolymerization of organic matter and nutrient acquisition involves also other enzymes, and therefore the enzyme investigated here may not fully represent C, N or P demand and availability (Sinsabaugh and Shah, 2012).

### Enzyme temperature sensitivity is similar through the soil profile across enzyme types

Our results confirmed the general expectation that activity of BG, LAP and AP is stimulated by warming, with Q_10_ values for *V*_max_ and activation energies within the ranges typically observed for soil exoenzymes (Brzostek and Finzi, 2012; German et al., 2012; Nottingham et al., 2016; Razavi et al., 2015, 2016; Steinweg et al., 2013; Stone et al., 2012; Trasar-Cepeda et al., 2007). However, despite few suggestive trends, temperature sensitivity did not vary significantly with depth for any enzyme or kinetic property based on either Q_10_, following a linear Arrhenius model, or *T*_opt_, TS_max_ and Δ*C*_p_^‡^ of *V*_max_ estimated by the non-linear MMRT model. We could not determine unambiguously whether the response of *V*_max_ to temperature was best explained by Arrhenius or MMRT models, although comparisons suggested that the latter generally fit the data better. Moreover, temperature response rates of *V*_max_ declined at higher temperatures in most cases, as observed by other studies (Alster et al., 2016a, 2018), further indicating that MMRT represented a more realistic temperature response behavior. On the other hand, this also indicated that the linear models captured the temperature response better under lower temperatures within the native temperature range, before response rates slowed towards *T*_opt_, similar to what has been observed by Alster et al. (2016a). The uniform temperature sensitivity observed here contradicted our initial hypothesis that exoenzymes in deeper soils are more sensitive to warming as a result of microbial adaptation to lower and narrower temperature ranges (Schipper et al., 2014). Previous studies have observed that exoenzymes from colder soil environments tend to be more sensitive to warming (Brzostek and Finzi, 2012; Koch et al., 2007), as well as in some subsoils in relation to surface soils (Steinweg et al., 2013). Therefore, we expected that Q_10_ values and activation energies would increase with depth, whereas either *T*_opt_, Δ*C*_p_^‡^, or both, would decrease. Under MMRT’s Optimum-Driven hypothesis, the more frequent lower temperatures in deeper soils could select for enzymes with lower *T*_opt_, regardless of their Δ*C*_p_^‡^ (Alster et al., 2020). Conversely, the Thermal Breadth hypothesis postulates that enzymes subject to large temperature ranges have less negative Δ*C*_p_^‡^ (i.e., flatter temperature response curves) but not necessarily different *T*_opt_, and thus the narrower temperature ranges in deeper soils would lead to more negative Δ*C*_p_^‡^ (Alster et al., 2020). In turn, the Enzyme Rigidity hypothesis predicts that cold-adapted enzymes have more negative Δ*C*_p_^‡^ due to their lower rigidity, which would lead to a decline in Δ*C*_p_^‡^, and consequently *T*_opt_, with depth, following selection of enzymes adapted to lower temperatures (Alster et al., 2020). The constant temperature sensitivity of all enzymes through the soil profile may instead reflect a convergence of enzyme *T*_opt_ towards the similar MATs across depths at our site (10.4-11.5°C), despite different temperature ranges. On the other hand, the low MATs of our soils appear to contradict the relatively high Δ*C*_p_^‡^ observed, compared to values previously reported (Alster et al., 2016a, 2018), which are expected to reflect a high enzyme rigidity typical of warm-adapted enzymes. It is possible that enzyme Δ*C*_p_^‡^ in our soils are mainly driven by their wide temperature ranges, despite their narrowing with depth, leading to selection of enzymes with less negative Δ*C*_p_^‡^, and thus able to maintain more constant activity rates under varying temperatures (Alster et al., 2020). Moreover, reactions potentially involving a diverse isozyme pool, such as those measured here, reflect the summation of the temperature response curves of those enzymes, and thus are also expected to have a less negative Δ*C*_p_^‡^ (Alster et al., 2018).

Our results are consistent with the uniform temperature sensitivity (apparent Q_10_) of *in situ* soil respiration over the top meter of soil previously observed at this site (Hicks Pries et al., 2017). This suggests that SOM depolymerization by exoenzymes may be closely linked to the response of total soil respiration to warming, likely by modulating the contribution of microbial heterotrophic metabolism. In turn, *K*_m_ was relatively insensitive to temperature, with mean Q_10_ values around 1 across depths and enzymes. This suggests that enzyme affinities may be biochemically constrained to prevent being affected by temperature fluctuations, as previously suggested (Allison et al., 2018). *T*_opt_ (65.19 ± 3.74°C) was much higher than natural soil temperatures, whereas TS_max_ (31.63 ± 1.98°C) was just above the temperature maximum in surface soils (29°C at 5 cm), but substantially higher than those at lower depths (19°C at 30 cm and 16°C at 100 cm), or mean annual temperatures (MAT) over the upper meter of soil (10.4-11.5°C). Similar high *T*_opt_ estimates based on MMRT have been generally observed for microbial exoenzyme activities and complex metabolic processes in soils (Schipper et al., 2014), and for soil bacterial isolates (Alster et al., 2016a). This is consistent with the fact that the thermal stability and optimal catalytic temperature of enzymes from mesophilic organisms tend to be higher than that of their native environment (Engqvist, 2018). While persistent warming is expected to induce higher enzyme activity through the whole soil profile, a uniform TS_max_ may, however, result in variable net annual temperature responses at different depths due to their different temperature ranges and seasonal variation. As response rates get faster when temperatures approach TS_max_, small temperature increases may disproportionally stimulate enzyme activity in surface soils in warmer months, when temperature frequently approaches TS_max_. However, warming above TS_max_ would cancel that effect. In turn, temperatures in deeper soils are never close to TS_max_, but sustained warming will bring them closer to it throughout the year within a narrower and more stable range than in surface soils, where temperature oscillates quicker, more frequently, and further away from TS_max_. How these possible scenarios may play out in response to sustained long-term warming will depend on the degree of thermal adaptability of exoenzymes through changes in microbial community composition, and expression of isozymes with different thermal properties (Bradford, 2013; Wallenstein et al., 2011). It should be noted, however, that we cannot completely rule out the possibility that the spatial variability of some temperature sensitivity estimates might have precluded detection of robust differences between depths.

### Different enzymes have similar temperature sensitivities but warming can affect their relative kinetics in a depth-dependent manner

The magnitude of all temperature sensitivity parameters was remarkably similar between enzymes through the soil profile, although Q_10_ values of *V*_max_ and *K*_m_ have been frequently shown to vary between co-occurring soil enzymes (Wallenstein et al., 2011). Likewise, *T*_opt_, TS_max_ and Δ*C*_p_^‡^ can vary substantially between enzymes (Alster et al., 2016a). Nevertheless, we did observe a significantly lower Q_10_ of *K*_m_ and consequently higher Q_10_ of CE of LAP in the upper 10 cm, relative to the other enzymes. The fact that the mean Q_10_ of *K*_m_ of LAP was below 1 (Q_10_ = 0.83), while those of BG and AP were not (Q_10_ = 1.30 and 1.18, respectively), suggested that the affinity of LAP was positively stimulated by warming at this depth (i.e., *K*_m_ decreased), or that those of BG and AP were negatively affected. As the Q_10_ of *V*_max_ did not differ between enzymes, this led to a higher positive temperature response of the CE of LAP. This likely contributed to the significant negative effect of higher temperatures on the CE^BG:LAP^ ratio, and shows that temperature can directly affect the relative catalytic efficiencies between C- and N-acquiring enzymes. This was consistent with previous observations suggesting that kinetic responses to temperature may vary among enzyme types, leading to changes in relative cycling of different nutrients (Allison et al., 2018). Moreover, the significant interaction between depth and temperature on CE^BG:LAP^ ratios confirmed that their variation with depth was dependent on temperature, possibly reflecting the higher CE Q_10_ of LAP in the upper 10 cm. Higher temperatures also had a significant positive effect on *V*_max_^BG:AP^ and *V*_max_^LAP:AP^ ratios, indicating that AP was generally less stimulated by warming than BG or LAP. As enzyme assays at different temperatures were performed with the same soil preparations per depth and incubated over relatively short periods, these relative differences in *V*_max_ likely reflected a direct effect of temperature on enzyme turnover (*K*_cat_), independently of enzyme concentration. Despite the uniform temperature sensitivity of individual enzymes through the soil profile, these results show that temperature can have different effects on intrinsic kinetic properties (i.e., *K*_m_ and *K*_cat_) of distinct enzymes in a depth-dependent manner, presumably without active microbial regulation.

## Conclusions

Kinetic and thermal properties of exoenzymes are fundamental components of complex trait spaces that allow microbes to thrive under variable nutrient availability and temperature regimes, as well as other interacting selective pressures (Allison et al., 2011; Ho et al., 2017; Malik et al., 2020; Ramin and Allison, 2019; Sinsabaugh et al., 2014; Sinsabaugh and Shah, 2012). Our results indicated that exoenzyme kinetics through the soil profile reflected, not only variation in substrate availability, but also different enzyme production levels and isozymes, which likely represent inherent traits of microbiomes with substrate and nutrient demands associated with distinct life strategies. Moreover, we show that the temperature sensitivity of exoenzymes is remarkably similar through the soil profile, but also that temperature can directly affect kinetic properties of different enzymes in a depth-dependent manner. This also implies that microbiomes at different depths produce isozymes with distinct thermal properties, which varies between enzyme types. Consequently, temperature can directly affect relative substrate depolymerization and nutrient acquisition potential, effectively decoupling enzyme relative activities from other regulatory factors, such as nutrient demand and substrate availability. Although microbial trait spaces are not static, as microbiomes adapt to changing conditions, they are likely to constrain both immediate responses and the trajectory of longer-term responses to environmental changes (Bradford, 2013; Conant et al., 2011; Xu et al., 2021). Therefore, it is essential to identify and validate key microbial traits and their environmental constraints in order to build a mechanistic understanding that can be generalized across spatiotemporal scales, and combine theory, measurements and models to improve the representation of microbial processes in Earth system models (Blankinship et al., 2018; Wieder et al., 2015). Together, our results improve the mechanistic understanding of microbial processes driving SOM dynamics as a function of soil depth and in response to warming, and provide new directions towards improved representation of key microbial traits in depth-resolved biogeochemical models.

## Data Availability Statement

Data and code generated for this study are available as Supplementary Materials, and can be found in the U.S. Department of Energy’s (DOE) Environmental System Science Data Infrastructure for a Virtual Ecosystem (ESS-DIVE) (Varadharajan et al., 2019) data repository [DOI provided upon publication].

## Author Contributions

RJEA, MM and ELB conceived and designed the study. RJEA, IAC, GM and ELB collected soil samples, and RJEA, IAC, HWS and BW performed soil chemical analyses and enzyme assays. RJEA analyzed and interpreted the data with support from IAC, GM, MM and ELB. RJEA wrote the manuscript with input from MM, MST and ELB. All authors read and reviewed the manuscript.

## Funding

This work was performed at Lawrence Berkeley National Laboratory and supported by the U.S. Department of Energy Office of Science, Office of Biological and Environmental Research under Contract No. DE-AC02-05CH11231 to LBNL as part of the Belowground Biogeochemistry Science Focus Area, through the Terrestrial Ecosystem Science Program. IAC was supported by a fellowship from the National GEM Consortium. HWS was supported by the University of California, Berkeley, Sponsored Projects for Undergraduate Research Program. BW was supported by the California Alliance for Minority Participation program sponsored by the National Science Foundation.

## Conflict of Interest

The authors declare that the research was conducted in the absence of any commercial or financial relationships that could be construed as a potential conflict of interest.

## Acknowledgements

We are grateful to Xiaoqin Wu and Romy Chakraborty (Lawrence Berkeley National Laboratory) for technical support with DOC and TDN measurements. We also thank William Riley, Jinyun Tang, and the whole Belowground Biogeochemistry Scientific Focus Area team at Lawrence Berkeley National Laboratory for helpful discussions and suggestions.

## Notes

### Competing Interest Statement

The authors have declared no competing interest.

