## Supplementary Information for "Kinetic Properties of Microbial Exoenzymes Vary with Soil Depth but Have Similar Temperature Sensitivities Through the Soil Profile"

### Supplementary Results

#### Comparison of temperature sensitivity based on Arrhenius and Macromolecular Rate Theory models *(extended)*.

Comparisons between alternative Arrhenius and MMRT models based on the AIC, AICc and BIC indices of model fit were largely inconclusive, although overall comparisons favored MMRT consistently by marginal differences. AICc consistently over-penalized the model with the highest number of parameters ($K$ = 3; MMRT) or the lowest number of temperature data-points ($n$ = 5; Arrhenius over 4-35°C), resulting in disproportionally high ΔAICc values in relation to the model with the lowest number of parameters and highest number of temperatures ($K$ = 2 and $n$ = 6; Arrhenius over 4-50°C) (Table S6). This was likely due to the low overall ratio between sample size (i.e., number of temperatures) and number of parameters ($n/K$ ratio) in our models, and the additional correction for this ratio used by AICc, which put a disproportional high weight on very small differences of just one parameter or data-point. Moreover, the number of parameters in our three-parameter MMRT model is effectively the same as that in the Arrhenius model ($K$ = 2), due to the interdependence between enthalpy and entropy parameters (Alster et al., 2020). The less stringent AIC index values were more consistent with the observed data trends, adjusted $R^2$ values of the linear models, and differences in $n/K$ ratio (Figures 5-6, Table S6). Absolute AIC values indicated that MMRT had the highest fit for 50% of the datasets, followed by Arrhenius over 4-50°C and then Arrhenius over 4-35°C. These results were mirrored by the BIC index, although this index has a greater penalty factor for low $n/K$ ratios than AIC (Tables S6). Nevertheless, ΔAIC values and AIC relative likelihoods showed substantial support for most alternative models, according to the guidelines by (Burnham and Anderson, 2004), indicating that, in general, the fit of alternative Arrhenius and MMRT models did not differ substantially (Tables S6). The few ΔAIC exceptions to this trend indicated that only 17% of the Arrhenius models over 4-50°C and 7% of those over 4-35°C had substantially worse fit than their two alternative models for the same dataset, whereas MMRT model fits were never substantially worse than their Arrhenius counterparts.

**Supplementary Tables**

**Table S1.** Soil and microbial biomass chemistry (available as spreadsheet in Additional_Supplementary_Tables.xlsx).

**Table S2.** Michaelis-Menten kinetic parameters (available as spreadsheet in Additional_Supplementary_Tables.xlsx).

**Table S3.** Two-way fixed effects ANOVA of kinetic parameters between enzymes per depth, with enzyme and temperature as independent factors. Differences were considered significant at $p < 0.05$.

| | | $V_{max/ds}$ | | | $V_{max/MBC}$ | | | $K_m$ | | | $CE_{ds}$ | | | $CE_{MBC}$ | | |
|---|---|---|---|---|---|---|---|---|---|---|---|---|---|---|---|---|
| **Depth (cm)** | **Factor** | **Df** | ***F* value** | ***p* value** | **Df** | ***F* value** | ***p* value** | **Df** | ***F* value** | ***p* value** | **Df** | ***F* value** | ***p* value** | **Df** | ***F* value** | ***p* value** |
| 00-10 | Enzyme | 2 | 359.48 | < 0.05 | 2 | 360.20 | < 0.05 | 2 | 246.78 | < 0.05 | 2 | 923.88 | < 0.05 | 2 | 2774.82 | < 0.05 |
| | Temperature | 5 | 30.50 | < 0.05 | 5 | 30.56 | < 0.05 | 5 | 1.98 | 0.11 | 5 | 26.31 | < 0.05 | 5 | 79.01 | < 0.05 |
| | Enzyme × Temperature | 10 | 0.22 | 0.99 | 10 | 0.22 | 0.99 | 10 | 5.04 | < 0.05 | 10 | 2.53 | 0.02 | 10 | 7.61 | < 0.05 |
| 10-20 | Enzyme | 2 | 61.57 | < 0.05 | 2 | 57.03 | < 0.05 | 2 | 57.62 | < 0.05 | 2 | 121.82 | < 0.05 | 2 | 28.54 | < 0.05 |
| | Temperature | 5 | 6.43 | < 0.05 | 5 | 6.43 | < 0.05 | 5 | 0.36 | 0.88 | 5 | 4.17 | 0.01 | 5 | 5.40 | < 0.05 |
| | Enzyme × Temperature | 10 | 0.32 | 0.97 | 10 | 0.95 | 0.50 | 10 | 0.49 | 0.885 | 10 | 0.17 | 1.00 | 10 | 1.37 | 0.24 |
| 30-40 | Enzyme | 2 | 114.38 | < 0.05 | 2 | 298.98 | < 0.05 | 2 | 128.68 | < 0.05 | 2 | 526.70 | < 0.05 | 2 | 697.14 | < 0.05 |
| | Temperature | 5 | 10.24 | < 0.05 | 5 | 26.77 | < 0.05 | 5 | 0.18 | 0.97 | 5 | 15.34 | < 0.05 | 5 | 20.31 | < 0.05 |
| | Enzyme × Temperature | 10 | 0.13 | 1.00 | 10 | 0.33 | 0.97 | 10 | 1.07 | 0.41 | 10 | 0.72 | 0.70 | 10 | 0.95 | 0.50 |
| 50-60 | Enzyme | 2 | 120.86 | < 0.05 | 2 | 207.69 | < 0.05 | 2 | 168.22 | < 0.05 | 2 | 302.36 | < 0.05 | 2 | 447.57 | < 0.05 |
| | Temperature | 5 | 11.44 | < 0.05 | 5 | 20.46 | < 0.05 | 5 | 1.00 | 0.43 | 5 | 9.49 | < 0.05 | 5 | 14.53 | < 0.05 |
| | Enzyme × Temperature | 10 | 0.39 | 0.94 | 10 | 0.59 | 0.81 | 10 | 0.78 | 0.65 | 10 | 0.63 | 0.78 | 10 | 0.88 | 0.56 |
| 60-70 | Enzyme | 2 | 76.55 | < 0.05 | 2 | 173.35 | < 0.05 | 2 | 85.51 | < 0.05 | 2 | 136.38 | < 0.05 | 2 | 276.20 | < 0.05 |
| | Temperature | 5 | 5.54 | < 0.05 | 5 | 12.58 | < 0.05 | 5 | 1.18 | 0.34 | 5 | 4.67 | 0.01 | 5 | 9.58 | < 0.05 |
| | Enzyme × Temperature | 10 | 0.31 | 0.98 | 10 | 0.68 | 0.73 | 10 | 0.87 | 0.57 | 10 | 0.10 | 1.00 | 10 | 0.14 | 1.00 |
| 80-90 | Enzyme | 2 | 145.88 | < 0.05 | 2 | 163.42 | < 0.05 | 2 | 30.63 | < 0.05 | 2 | 308.30 | < 0.05 | 2 | 345.72 | < 0.05 |
| | Temperature | 5 | 8.13 | < 0.05 | 5 | 9.50 | < 0.05 | 5 | 0.96 | 0.45 | 5 | 10.74 | < 0.05 | 5 | 12.34 | < 0.05 |
| | Enzyme × Temperature | 10 | 0.39 | 0.94 | 10 | 0.43 | 0.92 | 10 | 0.96 | 0.50 | 10 | 0.47 | 0.90 | 10 | 0.48 | 0.89 |

**Table S4.** Variation in Michaelis-Menten kinetic parameters between enzymes per depth. Different colors and letters indicate significant differences ($p < 0.05$), based on Tukey's tests after one-way ANOVA per depth (Table 2).

| Depth (cm) | Enzyme | $V_{max/ds}$ | $V_{max/MBC}$ | $K_m$ | $CE_{ds}$ | $CE_{MBC}$ |
|---|---|---|---|---|---|---|
| 00-10 | BG | b | b | b | a | b |
| | LAP | c | c | a | b | c |
| | AP | a | a | b | a | a |
| 10-20 | BG | b | b | c | a | a |
| | LAP | c | c | a | b | b |
| | AP | a | a | b | a | a |
| 30-40 | BG | b | b | c | a | a |
| | LAP | c | c | a | b | b |
| | AP | a | a | b | a | a |
| 50-60 | BG | b | b | c | b | b |
| | LAP | c | c | a | c | c |
| | AP | a | a | b | a | a |
| 60-70 | BG | b | b | b | b | b |
| | LAP | c | c | a | c | c |
| | AP | a | a | b | a | a |
| 80-90 | BG | b | b | b | b | b |
| | LAP | c | c | a | c | c |
| | AP | a | a | b | a | a |

**Table S5.** Two-way fixed effects ANOVA of ratios between kinetic parameters of BG, LAP and AP, with depth and temperature as independent factors. Differences were considered significant at $p < 0.05$.

| | $V_{max}^{BG:LAP}$ | | | $V_{max}^{BG:AP}$ | | | $V_{max}^{LAP:AP}$ | | |
|---|---|---|---|---|---|---|---|---|---|
| **Factor** | **Df** | ***F* value** | ***p* value** | **Df** | ***F* value** | ***p* value** | **Df** | ***F* value** | ***p* value** |
| Depth | 5 | 6.38 | < 0.05 | 5 | 24.81 | < 0.05 | 5 | 9.00 | < 0.05 |
| Temperature | 5 | 1.20 | 0.32 | 5 | 5.30 | < 0.05 | 5 | 2.41 | < 0.05 |
| Depth × Temperature | 25 | 0.35 | 1.00 | 25 | 0.82 | 0.70 | 25 | 0.35 | 1.00 |

| | $K_m^{BG:LAP}$ | | | $K_m^{BG:AP}$ | | | $K_m^{LAP:AP}$ | | |
|---|---|---|---|---|---|---|---|---|---|
| **Factor** | **Df** | ***F* value** | ***p* value** | **Df** | ***F* value** | ***p* value** | **Df** | ***F* value** | ***p* value** |
| Depth | 5 | 5.78 | < 0.05 | 5 | 11.32 | < 0.05 | 5 | 1.21 | 0.314 |
| Temperature | 5 | 2.14 | 0.07 | 5 | 2.32 | 0.05 | 5 | 0.50 | 0.77 |
| Depth × Temperature | 25 | 1.27 | 0.22 | 25 | 0.76 | 0.77 | 25 | 0.76 | 0.78 |

| | $CE^{BG:LAP}$ | | | $CE^{BG:AP}$ | | | $CE^{LAP:AP}$ | | |
|---|---|---|---|---|---|---|---|---|---|
| **Factor** | **Df** | ***F* value** | ***p* value** | **Df** | ***F* value** | ***p* value** | **Df** | ***F* value** | ***p* value** |
| Depth | 5 | 4.24 | < 0.05 | 5 | 15.91 | < 0.05 | 5 | 9.95 | < 0.05 |
| Temperature | 5 | 9.72 | < 0.05 | 5 | 0.63 | 0.68 | 5 | 2.21 | 0.06 |
| Depth × Temperature | 25 | 1.85 | < 0.05 | 25 | 0.59 | 0.93 | 25 | 0.26 | 1.00 |

**Table S6.** Fit indices for Arrhenius models over 4-50°C and 4-35°C, and MMRT model over 4-50°C (available as spreadsheet in Additional_Supplementary_Tables.xlsx).

**Table S7.** $Q_{10}$ coefficients and activation energy ($E_a$) estimated over 4-50°C and 4-35°C, and MMRT parameter estimates over 4-50°C from all individual models. Different background colors indicate different models or different temperature ranges. Values excluded from further analyses are indicated in red (see main text) (available as spreadsheet in Additional_Supplementary_Tables.xlsx).

**Table S8.** Mean $Q_{10}$ coefficients and activation energy ($E_a$) estimated over 4-50°C and 4-35°C, and MMRT parameter estimates over 4-50°C, per enzyme and depth. Different background colors indicate different models or different temperature ranges (available as spreadsheet in Additional_Supplementary_Tables.xlsx).

**Table S9.** One-way ANOVA of temperature sensitivity parameters between enzymes per depth. $Q_{10}$ coefficients were calculated over 4-35°C, and the MMRT model parameters $T_{opt}$, $TS_{max}$, and $\Delta C_p^{\ddagger}$ over 4-50°C. Differences were considered significant at $p < 0.05$.

| | $V_{max}$ $Q_{10}$ | | | $K_m$ $Q_{10}$ | | | CE $Q_{10}$ | | | $T_{opt}$ | | | $TS_{max}$ | | | $\Delta C_p^{\ddagger}$ | | |
|---|---|---|---|---|---|---|---|---|---|---|---|---|---|---|---|---|---|---|
| **Depth (cm)** | **Df** | ***F* value** | ***p* value** | **Df** | ***F* value** | ***p* value** | **Df** | ***F* value** | ***p* value** | **Df** | ***F* value** | ***p* value** | **Df** | ***F* value** | ***p* value** | **Df** | ***F* value** | ***p* value** |
| 00-10 | 2 | 0.28 | 0.76 | 2 | 14.58 | 0.01 | 2 | 35.90 | < 0.05 | 2 | 1.33 | 0.33 | 2 | 1.68 | 0.26 | 2 | 0.72 | 0.52 |
| 10-20 | 2 | 0.75 | 0.51 | 2 | 3.59 | 0.09 | 2 | 4.68 | 0.06 | 2 | 0.12 | 0.89 | 2 | 0.41 | 0.69 | 2 | 0.05 | 0.95 |
| 30-40 | 2 | 4.65 | 0.06 | 2 | 5.22 | 0.05 | 2 | 5.13 | 0.05 | 2 | 0.69 | 0.54 | 2 | 1.14 | 0.38 | 2 | < 0.01 | 1.00 |
| 50-60 | 2 | 3.80 | 0.09 | 2 | 0.39 | 0.69 | 2 | 1.98 | 0.22 | 2 | 0.30 | 0.75 | 2 | 0.19 | 0.83 | 2 | 1.10 | 0.40 |
| 60-70 | 2 | 6.38 | 0.03 | 2 | 1.60 | 0.28 | 2 | 0.05 | 0.95 | 2 | 4.00 | 0.09 | 2 | 2.00 | 0.23 | 2 | 3.66 | 0.11 |
| 80-90 | 2 | 1.16 | 0.38 | 2 | 0.21 | 0.82 | 2 | 0.12 | 0.89 | 2 | 2.38 | 0.17 | 2 | 1.98 | 0.22 | 2 | 1.09 | 0.40 |

**Table S10.** Variation in temperature sensitivity between enzymes per depth. $Q_{10}$ values were calculated over 4-35°C. Different colors and letters indicate significant differences ($p < 0.05$), based on Tukey's tests after one-way ANOVA per depth (Table S2).

| Depth (cm) | Enzyme | $V_{max}$ $Q_{10}$ | $K_m$ $Q_{10}$ | CE $Q_{10}$ |
|---|---|---|---|---|
| 00-10 | BG | a | a | b |
| | LAP | a | b | a |
| | AP | a | a | b |
| 10-20 | BG | a | a | a |
| | LAP | a | a | a |
| | AP | a | a | a |
| 30-40 | BG | a | a | b |
| | LAP | a | a | a |
| | AP | a | a | ab |
| 50-60 | BG | a | a | a |
| | LAP | a | a | a |
| | AP | a | a | a |
| 60-70 | BG | ab | a | a |
| | LAP | a | a | a |
| | AP | b | a | a |
| 80-90 | BG | a | a | a |
| | LAP | a | a | a |
| | AP | a | a | a |

**Supplementary Figures**

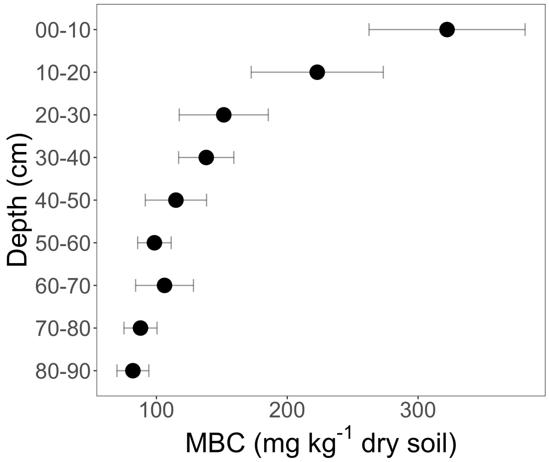


**Figure S1.** Microbial biomass carbon (MBC) concentration through the soil profile. Error bars indicate the standard error of the mean (n = 3).

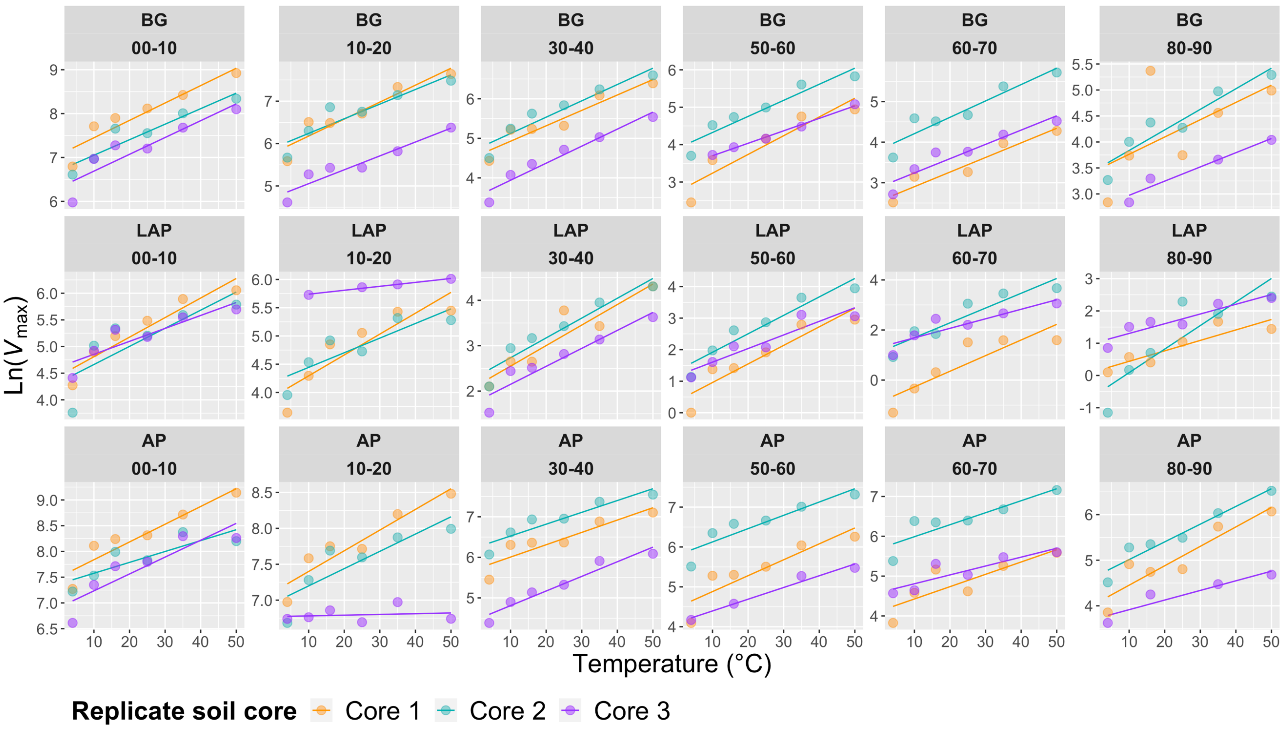


**Figure S2.** Arrhenius models of $V_{max}$ over five temperatures from 4 to 50°C per enzyme, depth and replicate core sample.

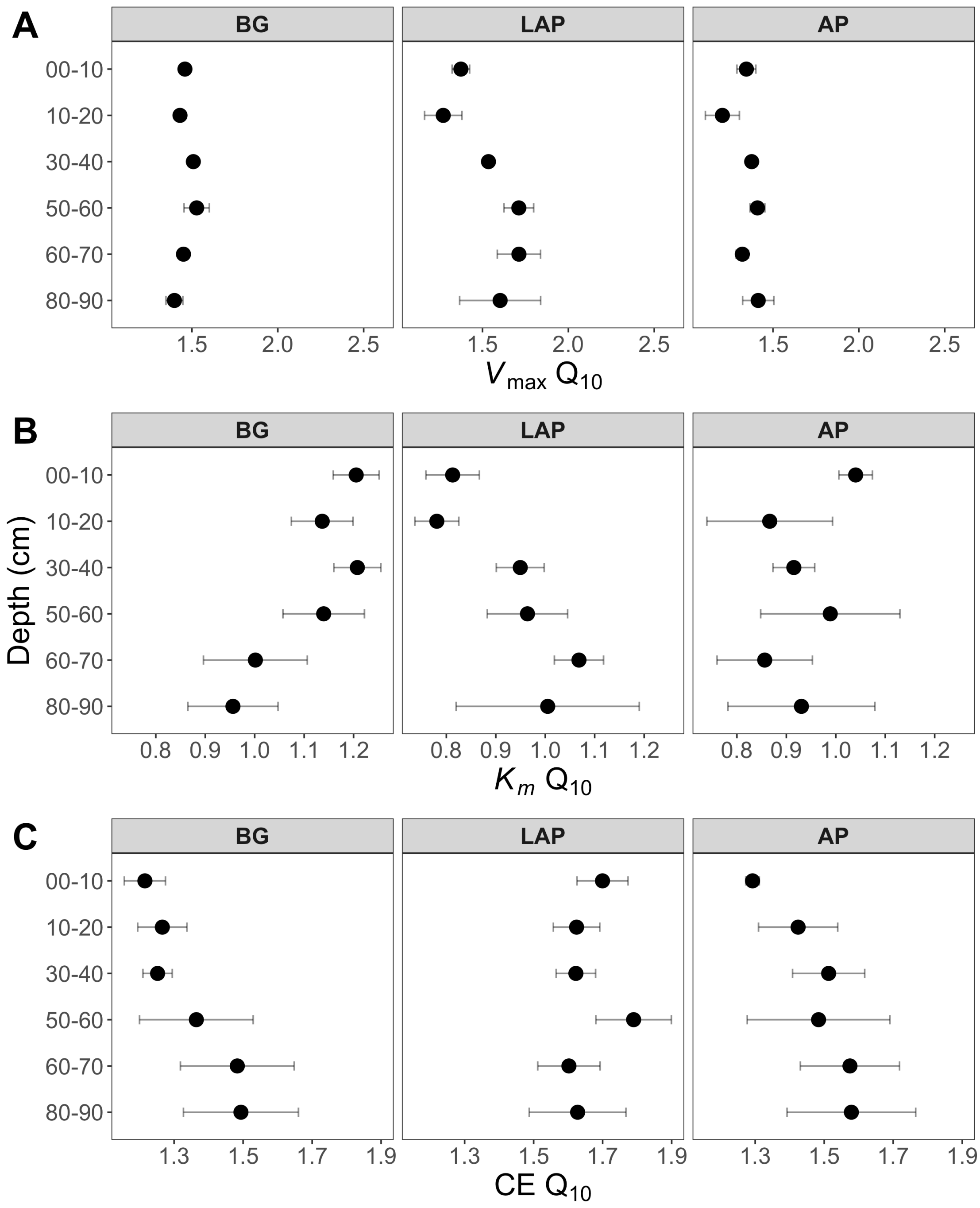


**Figure S3.** $Q_{10}$ coefficients of MM kinetic parameters over six temperatures from 4 to 50°C at different depths. (A) $V_{max}$, (B) $K_m$, and (C) CE. Differences between depths are not significant ($p > 0.05$), based on ANOVA per enzyme (Table 2). Error bars represent the standard error of the mean (n = 3).

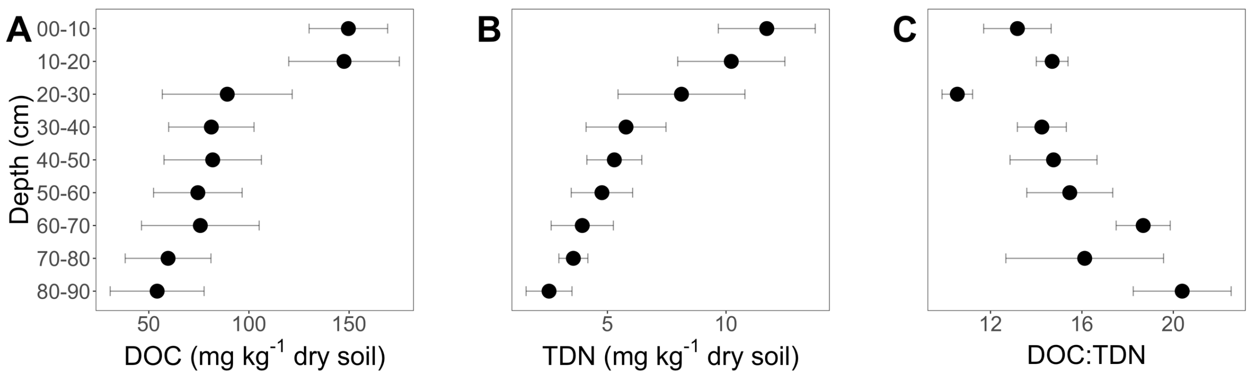


**Figure S4.** Soil (A) dissolved organic carbon (DOC), (B) total dissolved nitrogen (TDN), and (C) DOC:TDN ratio. Error bars indicate the standard error of the mean (n = 3).
